# Thrombospondin-2 deficiency primes the synovial joint for aberrant tissue remodeling and injury response

**DOI:** 10.64898/2026.06.18.732872

**Authors:** Lindsey Lammlin, Lucas M. Junginger, Alexander J. Knights, Michael D. Newton, Henry Dai, Carlisle R. DeJulius, Aanya Mohan, Isabelle J. Smith, Scarlet C. Howser, Gurjit S. Mandair, Sunghun Cheong, Paul F. Lais, Sofia Gonzalez-Nolde, Andrea I. Alford, Kurt D. Hankenson, Tristan Maerz

## Abstract

**Objective:** This study investigates joint injury-induced angiogenesis and the effects of genetic deficiency of thrombospondin-2 (TSP2), an anti-angiogenic factor, in joint homeostasis and post-traumatic osteoarthritis (PTOA).

**Method:** We utilized a murine non-invasive anterior cruciate ligament rupture (ACLR) model of PTOA and mined published synovial transcriptomics datasets to investigate injury-induced synovial angiogenesis. Spatial transcriptomics and flow cytometry of TSP2-GFP reporter mice were used to assess injury-induced thrombospondin-2 and its cellular origins in synovium. Global TSP2 knockout mice (TSP2-KO) were used to assess the effect of TSP2 deficiency on early and late stages of PTOA development via molecular imaging of inflammation and angiogenesis, histopathology, micro-computed tomography, Raman spectroscopy, and synovium bulk RNA-sequencing.

**Results:** Intra-articular angiogenesis peaked at 7d post-ACLR and declined but remained elevated above baseline at 28d post-ACLR. We identified synovial crosstalk between endothelial cells and sublining fibroblasts as a key driver of angiogenesis and source of thrombospondin-2 signaling, with TSP2 primarily upregulated in sublining fibroblasts. TSP2-KO mice exhibited increased peri-articular inflammation at 7d post-ACLR and inferior bone quality. Histopathology revealed greater PTOA severity but paradoxically lower synovitis in TSP2-KOs. Additionally, aberrant structural remodeling of the entire knee joint was observed in uninjured and ACLR TSP2-KO limbs. The uninjured TSP2-KO synovial transcriptome demonstrated elevated immune, fibrotic, and angiogenic activation; however, TSP2-KO and WT synovial transcriptomes partially converged upon injury.

**Conclusion:** TSP2 is essential for joint homeostasis and trauma response. Global TSP2 deficiency causes premature OA and worsened PTOA, suggesting that therapeutic targeting with TSP2 mimetic could be used to prevent OA.

## INTRODUCTION

Intra-articular angiogenesis is at the center of the joint’s inflammatory response to trauma and is a driver of the development of chronic synovitis during post-traumatic osteoarthritis (PTOA). Joint injury activates angiogenesis in the synovium, increasing vascular access of immune cells to synovium to perpetuate synovitis^1,2^. Blood vessel infiltration is also observed in normally avascular articular cartilage and menisci, and this angiogenesis has been linked to the downstream activation of pathological processes such as chondrocyte hypertrophy, ectopic mineralization of cartilage, and osteophyte formation^1,3–7^. Existing clinical evidence indicates an association between synovial blood vessel density and worse OA symptoms and severity^1,8^.

The cellular and molecular events that promote trauma-induced angiogenesis in the joint require extensive additional study. Prior studies demonstrated that following joint injury, activated synovial macrophages, synovial fibroblasts, and chondrocytes release pro-angiogenic factors to promote recruitment and activation of vasculogenic cells ^6,9–11^. Vascular endothelial growth factor (VEGF) has been described as a critical pro-angiogenic protein expressed by multiple cell types following injury^10,12,13^. Therapeutic targeting of VEGF has shown efficacy in preclinical models of collagen-induced arthritis^14,15^ and PTOA^16^, and prior studies employing murine PTOA models have shown that blocking angiogenesis at the time of injury mitigates disease development^17,18^.

As most research has focused on positive regulators of angiogenesis, studies of negative angiogenic mediators are lacking in OA. The thrombospondins are a family of five secreted matricellular proteins of which the structurally similar thrombospondin-1 (TSP1/*THBS1*) and thrombospondin-2 (TSP2/*THBS2*) have described anti-angiogenic functions^19–21^. Focusing on TSP2, its inhibitory effects on angiogenesis have been described by several mechanisms, including suppressing MMP2 and MMP9 activity, which limits VEGF bioavailability^22,23^ as demonstrated in murine wound healing and malignant glioma models^24,25^. TSP2 also interacts directly with endothelial cell integrins, CD36, and CD47, to impair migration, proliferation, and survival^26–29^. Concordant with TSP2’s role in modulating angiogenesis, TSP2-KO mice exhibit increased vascularity of the skin, adipose, thymus, retina, and bone^30–34^. Beyond its anti-angiogenic functions, TSP2 has also been implicated in the regulation of extracellular matrix (ECM) assembly^35^, and studies in TSP2-deficient mice demonstrate abnormalities in collagen fibrillogenesis and matrix ultrastructure^36–38^. Mechanistically, TSP2 influences ECM assembly through modulation of MMP activity and collagen cross-linking, in addition to regulating cell–matrix interactions^35^. Limited evidence exists regarding the role of TSP2 in joint homeostasis and disease. In human rheumatoid arthritis (RA) synovial tissue, TSP2 expression was inversely correlated with synovial angiogenesis^20^. In a chimera of human synovium implanted in severe combined immunodeficiency (SCID) mice, intraperitoneal treatment with TSP2-transfected fibroblasts limited vascular invasion of the implanted human synovium^20^.

Despite prior evidence demonstrating the functional effects of TSP2 in the regulation of tissue vascularity and matrix organization, its role in joint injury and PTOA pathogenesis has not been assessed. We discovered TSP2 as a highly induced factor in the synovium of injured mice, and here we show that TSP2-knockout mice exhibit aberrant idiopathic joint remodeling and exacerbated joint injury responses, establishing TSP2 as an important regulator of joint homeostasis and PTOA pathogenesis.

## MATERIALS AND METHODS

Expanded methods for each section is provided in the **Supplemental Methods**.

### Mice and PTOA model

All animal studies were approved by the University of Michigan Institutional Animal Care and Use Committee. Mice of three different genotypes were used: C57Bl/6J mice (wildtype/WT, Jackson Laboratories #000664), global thrombospondin-2 null mice (TSP2-KO, Jackson Laboratories #006238)^32^, and TSP2-GFP reporter mice^39^. Male and female mice aged 12-14 wks were used for joint injury studies. Post-traumatic osteoarthritis was induced via non-invasive anterior cruciate ligament rupture (ACLR) as described previously^40^ (**Suppl.-Methods**). Phenotyping of naïve, mature adult mice occurred in WT and TSP2-KO females aged 36-42 wks.

### Near-infrared (NIR) imaging

AngioSense 750 (Revvity Cat. #NEV10011EX) was used to assess intra-articular angiogenesis in WT mice at 7d and 28d post-ACLR. A separate cohort of WT and KO mice was imaged at 7d post-ACLR using both AngioSense 750 and the pan-cathepsin activatable probe ProSense 680 (Revvity Cat. #NEV10003) to assess genotypic differences in inflammation, angiogenesis, and vascular permeability. All imaging was conducted 4 hours post-injection, and signal quantification was conducted in a blinded fashion in FIJI/ImageJ (**Suppl.-Methods**).

### Mining of publicly available transcriptomics datasets

To assess genes and pathways related to synovial angiogenesis, we mined our previously published mouse synovium bulk RNA-seq^40^ and single-cell RNAseq (scRNAseq)^41^ data, comprised of synovium from both sexes across healthy (Sham), 7d ACLR, and 28d ACLR mice. We extracted endothelial cells from scRNAseq data to examine the injury-induced gene expression changes at 7d and 28d post-ACLR relative to Sham. Differentially expressed genes (*P_adj_* < 0.05) within the “Angiogenesis” GO:BP term at 7d or 28d post-ACLR relative to Sham were identified by bulk RNA-seq and mapped onto our synovial scRNAseq clusters to assess cell type-specific expression. CellChat was used to infer cellular communication networks by cell type and identify ligand-receptor interactions. scRNAseq data was also mined to identify cell type-specific expression patterns of the thrombospondin gene family and the integrin receptors *Cd36* and *Cd47* (**Suppl.-Methods**).

### Flow cytometry

Synovia from TSP2-GFP reporter mice were harvested 3d post-ACLR, digested to yield single cells, and stained with various antibodies to identify TSP2-GFP-expressing cells by flow cytometry using a BD LSR Fortessa (**Suppl.-Methods**, **Suppl.-Table 1**).

### Cryohistology

TSP2-GFP reporter limbs were harvested 3d, 14d, and 28d post-ACLR, fixed, decalcified, and processed for cryohistology. Sagittal joint tissue sections were stained with DAPI and imaged via fluorescent microscopy. TSP2-GFP+ synovial cells were quantified as a percentage of all synovial cells (**Suppl.-Methods**).

### Synovial tissue gene expression

Paired contralateral and ACLR synovia were harvested 3d and 14d post-ACLR into TRIzol and stored at −80°C until analysis. Samples were thawed, homogenized, and RNA was isolated via chloroform precipitation. cDNA synthesis was performed followed by RT-qPCR of *Thbs2*, with *Atp5b* as a housekeeper (**Suppl.-Methods**).

### In vitro experiments

Primary fibroblast-like synoviocytes (FLS) were isolated from hindpaws of naïve adult male and female C57Bl/6 mice, as previously described^41^. Passage 3-4 cells were used in all experiments. FLS were seeded in 12 well plates, serum starved overnight, then exposed to serum-free media containing PBS (control), TNF-α (10 ng/mL), Wnt3a (10 ng/mL), or TGF-β (10 ng/mL) for 24 hours. Following treatment, cells were lysed with TRIzol and stored at −80**°**C, followed by RNA isolation and gene expression studies by RT-qPCR.

### Xenium Prime 5K spatial transcriptomics

Hindlimbs from Sham, 7d ACLR, and 28d ACLR WT mice (n=2 per sex/group) were collected for spatial transcriptomics according to our published protocol^42^ (**Suppl.-Methods**). Briefly, following cardiac perfusion, hindlimbs were removed, fixed, decalcified, and paraffin processed. Samples were individually embedded, trimmed, and RNA quality was assessed for each sample. Individual samples were re-embedded into a single block, from which a section was processed using the Xenium Prime 5K platform. Synovial tissues were manually outlined in Xenium Explorer and major cell populations within the synovium were annotated. Transcripts of interest were overlayed onto the cell annotations and visualized in Xenium Explorer.

### Raman spectroscopy

Hindlimbs of naïve mature adult KO and age-matched WT mice were harvested, similarly fixed (**Suppl. Methods–Cryohistology**), and processed for plastic-embedded confocal Raman spectroscopy^43^. Standard bone mineralization and matrix quality parameters were assessed in epiphyseal trabecular bone and subchondral bone of the femur (n=2 per sex/genotype)^44^.

### Micro-computed tomography (μCT)

Hindlimbs were formalin-fixed and then stored in 70% EtOH until ready for µCT scanning (SkyScan 1176; 8.9-µm voxel). Femoral bone morphometry was evaluated by compartment using a neural net segmentation algorithm and osteophytes were manually contoured and quantified. Dimensionality reduction by sPLSDA was performed using MetaboAnalyst^40^ (**Suppl.-Methods**).

### Paraffin histology and histopathological scoring

Fixed limbs were decalcified, paraffin processed, embedded in paraffin, sectioned (5 µm), and stained using Safranin-Orange and Fast Green. Sections were imaged (10X, Eclipse Ni E800 with DS-Ri2 camera, Nikon, Tokyo, Japan) and evaluated for synovitis severity (**Suppl.-Table 2**) and PTOA severity (**Suppl.-Table 3**) by a blinded scorer (n=3-12 sections were scored and averaged together per limb).

### Bulk RNA-sequencing

Synovia, inclusive of Hoffa’s fat pad, were harvested from WT and KO Sham, 7d ACLR and 28d ACLR mice (n=3 per sex/genotype/injury group) for RNAseq (**Suppl.-Methods**). Bioinformatic analysis was performed in RStudio (R v4.4.0, RStudio v2024.04.1). Quality control and principal component analysis (PCA)-based outlier analysis was performed. Differential gene expression analysis via weighted Limma-Voom compared genotype differences across Sham, 7d ACLR, and 28d ACLR groups. Pathway analysis was performed using gene set enrichment analysis (GSEA), statistical over-representation analysis, and Reactome analysis, employing the Gene Ontology-Biological Processes (GO:BP) and Reactome Pathway annotations (**Suppl.-Methods**).

*Statistical analysis:*

SPSS (v30, IBM) and Prism 9.0 (Graphpad) were used for statistical analyses which included multiple paired and not paired t-tests, and two-way linear mixed-effects models with Sidak post-hoc correction where appropriate (**Suppl.-Methods**).

## RESULTS

### Joint injury induces synovial angiogenesis

Employing the noninvasive ACLR model in C57BL/6 mice, we first characterized the synovial angiogenic response to injury. In this model, injury triggers a robust synovial immune response in the first 1-3 days^45^, followed by expansion and phenotypic diversification of synovial fibroblasts by 7d post-ACLR^41^. Following this acute synovitis, mice develop progressive PTOA and transition to chronic synovitis up to 28 days post-ACLR^40^. We performed intravital near-infrared imaging using the AngioSense 750 probe to assess joint vascularity and observed that ACLR joints exhibited higher AngioSense signal compared to uninjured/contralateral joints during inflammatory (7d) and established disease (28d) stages (Fig.-1A-B). Quantifying the angiogenesis-relevant expansion of endothelial cells, flow cytometry demonstrated a 7.3-fold increase in total CD31(+) endothelial cells in ACLR relative to contralateral synovia (**Fig.-1C**) at 3d post-ACLR, preceding the increased vascular density measured at 7d post-ACLR.

**Figure 1.**
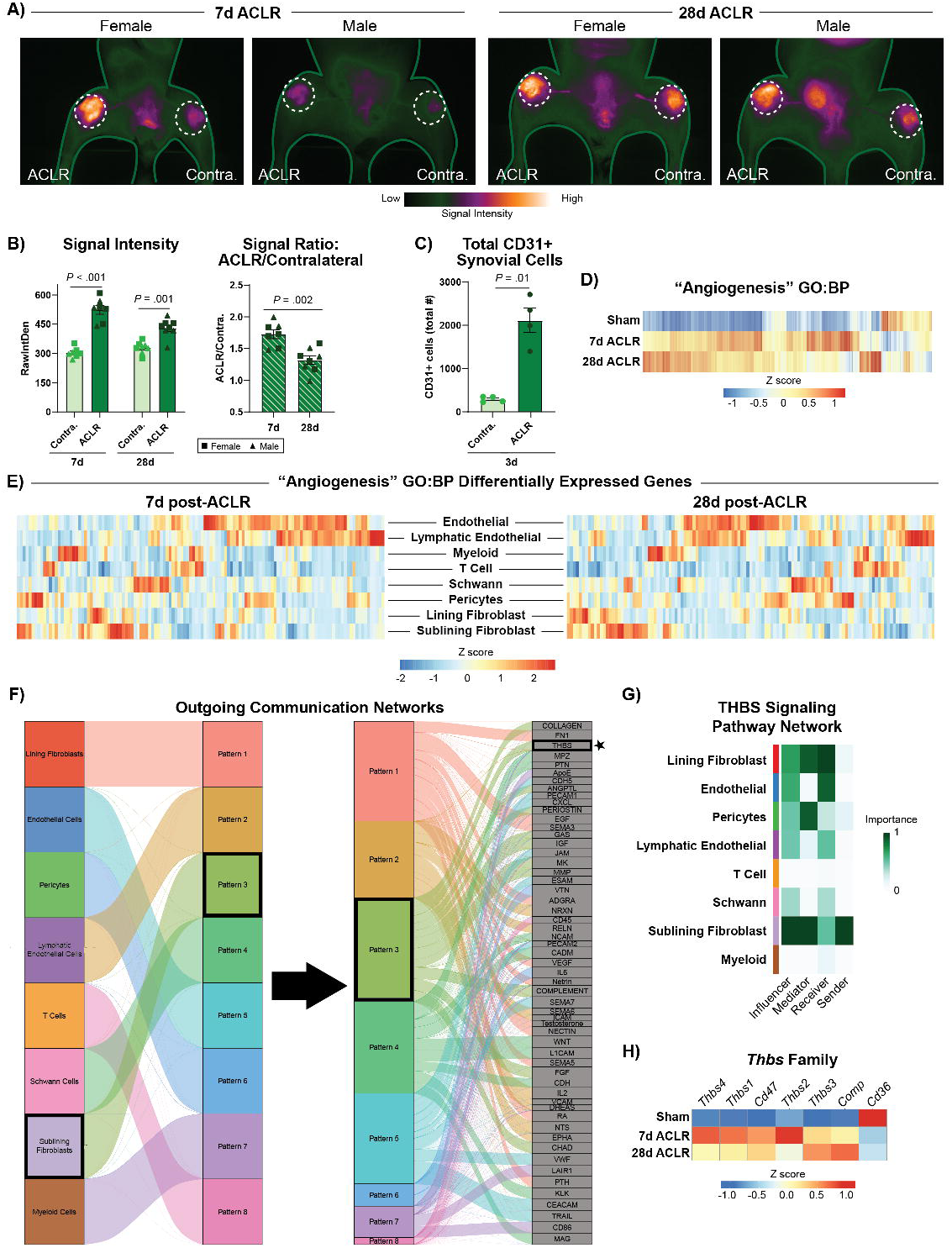
Intra-articular angiogenesis following joint injury. **A)** Representative intravital fluorescence imaging heatmap of AngioSense 750 at 7d and 28d post-ACLR. **B)** NIR signal intensity quantification of raw integrated density (RID) in ACLR limbs and the ACLR/Contralateral RID signal ratio (n=7-8 per timepoint; paired t-test between contralateral and ACLR limbs within each timepoint; t-test between timepoints). **C)** Total number of CD31+ cells in contralateral and ACLR synovia 3d post-ACLR (n=4; paired t-test between contralateral and ACLR synovia. All bars show mean ± SEM. **D)** Heatmap of genes within the “Angiogenesis” Gene Ontology (GO:0001525) term across healthy (Sham), 7d, and 28d post-ACLR osteoarthritic mouse synovial bulk RNAseq data^40^. **E)** Differentially expressed genes within the “Angiogenesis” GO term shown in **(D)** at 7d or 28d post-ACLR relative to Sham mapped across synovial cell types from published osteoarthritic mouse single cell RNAseq data^41^. **F)** Cell chat cellular communication analysis showing outgoing communication patterns across synovial cell types. Pattern 3, which originates from sublining fibroblasts is highlighted along with its constituent signaling term, THBS. **G)** THBS signaling pathway network across synovial cell types. **H)** Expression of the thrombospondin family of genes and the thrombospondin receptors *Cd36* and *Cd47* in mouse synovium bulk RNAseq^40^.

To characterize the molecular programs underpinning injury-induced synovial angiogenesis, we re-analyzed our published mouse bulk RNAseq data from male and female Sham, 7d ACLR, and 28d ACLR synovia^40^. This demonstrated substantial injury-induced perturbation to the “Angiogenesis” Gene Ontology – Biological Processes (GO:BP) pathway term at both 7d and 28d post-ACLR (**Fig.-1D**), showing activation of angiogenic processes in both early and late stages of PTOA. Significant DEGs in this term were mapped onto single-cell RNAseq from our published atlas of murine synovium^41^, which highlighted endothelial cells, lymphatic endothelial cells, and sublining fibroblasts as key contributors to injury-induced angiogenesis (**Fig.-1E**). Focusing on specifically synovial endothelial cells, top genes upregulated at 7d post-injury included the proinflammatory cytokine *Il6*, known positive regulators of endothelial cell activation and vascular sprouting (*Mmp14, Rcan1, Fam167b)*) and multiple collagen genes (*Col1a1/Col1a2, Col3a1, Col18a1, Col5a2*) (**Suppl.-Fig.-1A**). Collagen genes were abundant in the top DEGs at 28d post-injury, but negative regulators of angiogenesis were also upregulated (e.g. *Gadd45g*), suggesting chronic remodeling. Top downregulated genes after injury included negative regulators of angiogenesis (*Timp4, Car4)* and genes involved in the regulation of cellular metabolism (*Xdh, Atp5md*).

Leveraging scRNAseq data^41^, we next examined angiogenesis-relevant cellular communication and signaling networks using CellChat. We focused on outgoing communication from sublining fibroblasts as they represented the major non-endothelial source of angiogenesis-related DEGs (**Fig.-1E**). Sublining fibroblasts were predicted to send multiple angiogenesis-related signals, including known positive regulators such as VEGF, FGF, and ANGPTL. Focusing on negative regulators of angiogenesis, we identified thrombospondin (THBS) signaling as a predicted sublining fibroblast-derived communication axis (**Fig.-1F**). The THBS signaling network shows sublining fibroblasts to be the strongest senders while endothelial cells and lining fibroblasts were the strongest predicted receivers of thrombospondin signals (**Fig.-1G**). In RNAseq data of murine synovium^40^, we identified robust injury-induced upregulation of the entire thrombospondin family, with *Thbs2* (the gene encoding TSP2) exhibiting the greatest injury-induced upregulation at 7d post-ACLR (**Fig.-1H**), coinciding with peak vascularity measured by AngioSense. We confirmed *Thbs2* upregulation at additional timepoints via RT-qPCR (**Suppl.-Fig.-2A**). Assessing ligand-receptor contributions in the THBS signaling network, we observed substantial contributions from the interactions between TSP2 and its primary receptors CD36 and CD47 (**Suppl.-Fig.-1B**). In bulk RNA-seq, *Cd47* was modestly upregulated at 7d and 28d (**Fig.-1H**). Despite the large injury-induced increase in synovial endothelial cells (**Fig.-1C**), *Cd36* was downregulated at both timepoints (**Fig.-1H**). Taken together, these findings demonstrate that joint injury causes robust activation of synovial angiogenic processes, including thrombospondin signaling between sublining fibroblasts and endothelial cells.

### Sublining synovial fibroblasts activate thrombospondin-2 expression following joint injury

To spatially contextualize the observed induction of TSP2, we employed TSP2-GFP transgenic reporter mice, which express GFP driven by the endogenous *Thbs2* promoter^39^. Fluorescent cryosections demonstrated a near absence of TSP2-GFP+ cells in the uninjured/contralateral joint, besides expected TSP2-GFP in periosteum ^46^ (Fig.-2A). Joint injury induced a large expansion of TSP2-GFP+ cells in the synovium as early as 3d post-ACLR, with sustained expression at 14d and 28d post-ACLR during PTOA progression (**Suppl.-Fig.-2B**). Spatially, TSP2-GFP+ cells were confined to the synovial sublining and infrapatellar fat pad while the hyperplastic synovial lining was devoid of TSP2-GFP+ cells (**Fig.-2A**).

**Figure 2.**
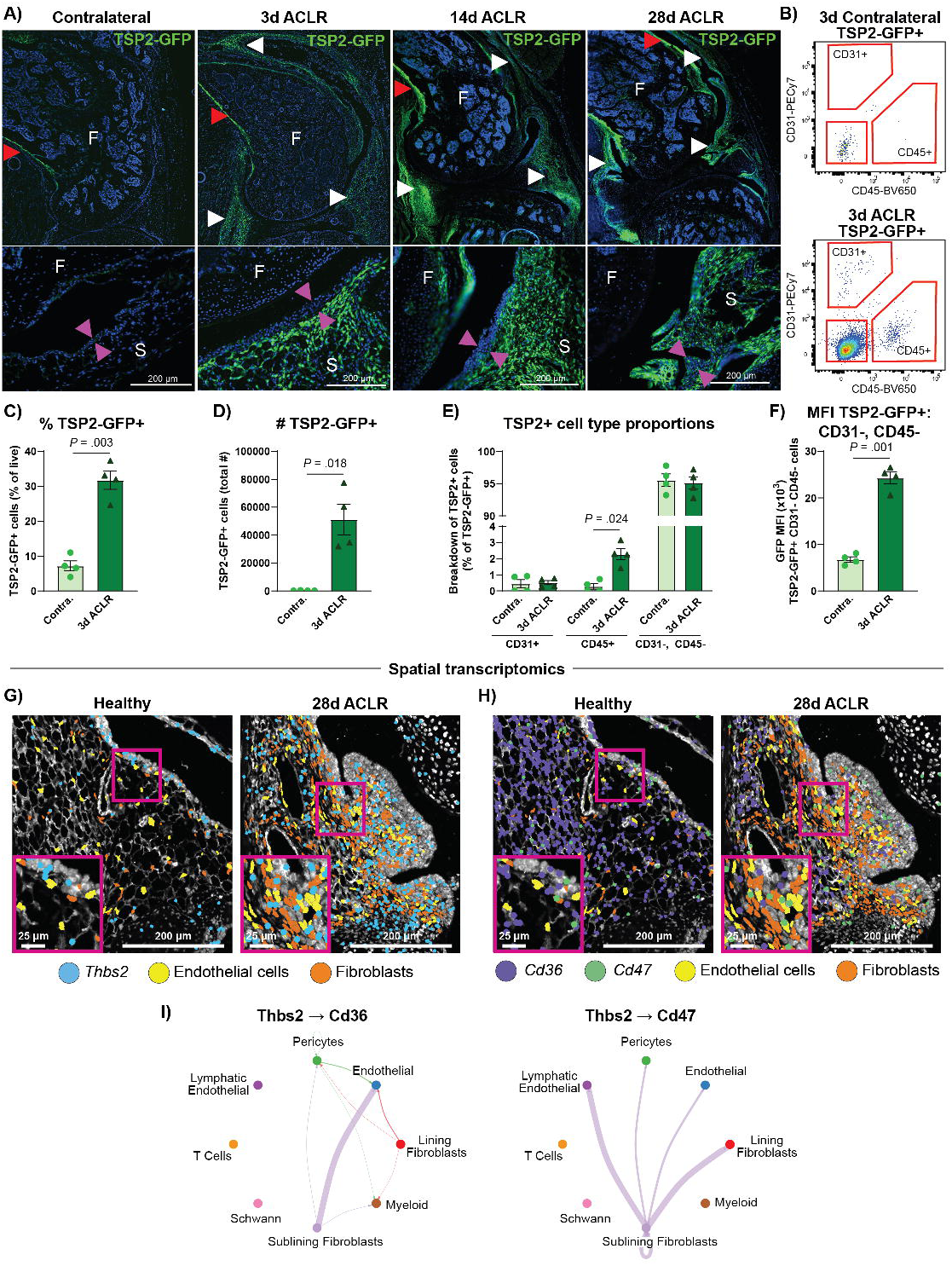
TSP2 induction in synovial cells, primarily fibroblasts, increases in abundance after joint injury. **A**) Representative uninjured contralateral, 3d-, 14d-, and 28d ACLR images of TSP2-GFP reporter limbs with DAPI nuclear stain (Red arrows: TSP2-eGFP+ bone. White arrows: TSP2-GFP+ synovium. Pink arrows: synovial lining layer). **B-F)** TSP2-GFP reporter mice were subjected to ACLR and synovia were harvested for flow cytometry 3d post-ACLR (n=4 mice). **B)** Representative plots of TSP2-GFP+ cell-type gating. **C)** Percentage of live cells that were TSP2-GFP+ in contralateral and 3d ACLR synovium. **D)** The total number TSP2-GFP+ cells in contralateral and 3d ACLR synovium. **E)** The breakdown of TSP2-GFP+ cells into CD31+ endothelial cells, CD45+ hematopoeitic cells, or CD31-CD45-synovial fibroblasts, in contralateral or 3d ACLR synovium. Paired two-tailed student’s t-tests were performed to assess significance in (B-F). **F)** Median fluorescence intensity (MFI) of TSP2-GFP in CD31-CD45-cells from contralateral or 3d ACLR synovium. All bars show mean ± SEM. **G-H)** Xenium 5k spatial transcriptomics gene overlays of **(G)** *Thbs2* (blue) or **(H)** the TSP2 receptors, *Cd36* (purple) and *Cd47* (green) with cell annotations (endothelial cells – yellow, sublining fibroblasts – orange, pink – zoom boxes). **I)** Circle plots showing intercellular communication between TSP2 ligand-receptor pairs.

To identify TSP2-expressing cells, we performed flow cytometry of whole synovium from TSP2-GFP mice at 3d post-ACLR (**Fig.-2B, Suppl.-Fig.-3A-B**). Injured synovia had a large increase in the number and proportion of TSP2-GFP+ cells relative to contralateral at 3d post-ACLR (**Fig.-2C-D**). TSP2-GFP+ cells in injured synovia were primarily stromal cells (CD31-, CD45-; 95% of total) and CD31(-), CD45(+) immune cells (2% of total) (**Fig.-2E**). Although CD45(+) CD11b(+) CD3(-) myeloid cells represented a small proportion of the TSP2-GFP+ cells, their number increased significantly following ACLR (**Suppl.-Fig.-3C-E**). The GFP median fluorescence intensity of TSP2-GFP-expressing stromal cells and all TSP2-GFP-expressing cells was increased in ACLR versus contralateral synovia (**Fig.-2F, Suppl.-Fig.-3F**), indicating injury-induced activation of *Thbs2* gene expression. We corroborated these findings using scRNAseq data, demonstrating that *Thbs2* is highly expressed by *Prg4*-Lo *Thy1*(+) *Pdgfra*(+) sublining fibroblasts. Lower but detectable expression was observed in a subset of pericytes, myeloid cells, and endothelial cells (**Suppl.-Fig.-2C**). Focusing on the TSP2 receptors, *Cd36* was highly expressed by endothelial cells and a subset of myeloid cells and pericytes, and *Cd47* was expressed ubiquitously (**Suppl.-Fig.-2C-D**).

We next utilized Xenium Prime 5K spatial transcriptomics, a probe-based assay with single-cell resolution, to visualize the spatial distribution of *Thbs2*, *Cd36*, and *Cd47* transcripts and their relevant cell types in synovium and infrapatellar fat pad of healthy and ACLR 28d murine joints. This corroborated the injury-induced enrichment of *Thbs2* and confirmed its overwhelming localization to the synovial sublining (**Fig.-2G-H**). This also corroborated the decrease in *Cd36* transcript despite the robust enrichment of endothelial cells, which may be indicative of dysfunctional vascularity recently associated with synovial damage and patient-reported symptoms in human OA^47^. Using scRNAseq data^41^, we inferred *Thbs2* cell type-specific communication networks, which showed that Thbs2-to-Cd36 signaling primarily originated in sublining fibroblasts and terminated in endothelial cells, whereas the Thbs2-to-Cd47 axis primarily signaled from sublining fibroblasts to lymphatic endothelial cells and lining fibroblasts (**Fig.-2I**).

Having identified synovial fibroblasts as the primary source of TSP2/*Thbs2*, we sought to identify factors that induce *Thbs2* expression in primary cultured synovial fibroblasts *in vitro*. The Wnt ligand, WNT3a, and TGF-β, both of which are fibrosis-relevant mediators^48^ that are upregulated in the synovium following ACLR, activated *Thbs2* expression (**Suppl.-Fig.-2E**). The pro-inflammatory cytokine TNF-α suppressed *Thbs2* (**Suppl.-Fig.-2E**).

Taken together, these results demonstrate that sublining synovial fibroblasts underpin the rapid and sustained upregulation of TSP2 expression following joint injury.

### TSP2-KO mice have greater injury-induced cathepsin protease activity and are predisposed to early PTOA development

Given the canonical anti-angiogenic function of TSP2, we evaluated peri-articular inflammation and vascular density in wildtype (WT) and TSP2 knockout (KO) mice at 7d post-ACLR using the molecular imaging probes ProSense 680, a cathepsin-activated marker of broad inflammation, and AngioSense 750. KOs exhibited greater ProSense 680 signal in the 7d ACLR limb while contralateral signal and ACLR/contralateral signal ratios were similar across genotypes (Fig.-3A-B**, Suppl.-Fig.-4A**). AngioSense showed the expected ACLR-induced signal increase with no genotype differences (**Suppl.-Fig.-4B-C**). This suggests that KOs exhibit greater injury-induced inflammation but similar vascular density/permeability as WTs.

**Figure 3.**
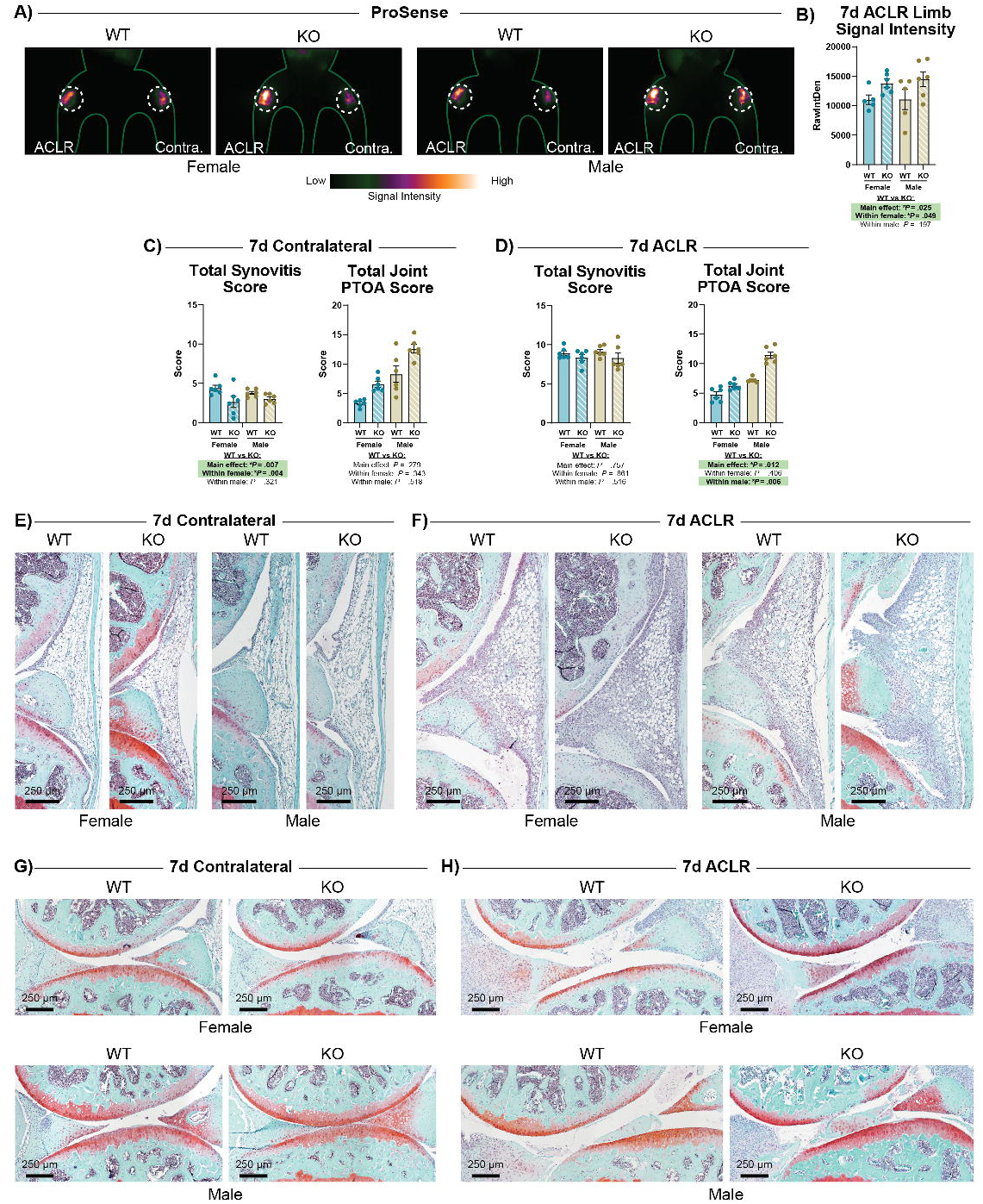
TSP2 KO vs WT 7d post-ACLR inflammation, angiogenesis, and structural pathology. **A)** Representative intravital fluorescence imaging heatmap of ProSense 680 at 7d post-ACLR. **B)** ProSense 680 NIR signal quantification in the 7d ACLR limb, expressed as raw integrated density (RID) (n=5-6 per sex/genotype). Histopathologic scoring of Safranin-Orange and Fast Green stained joints to evaluate synovitis severity and PTOA severity in the **(C)** contralateral and **(D)** 7d ACLR limb. The synovitis score evaluates the anterior synovium compartment while the PTOA score represents a composite score of the entire tibiofemoral joint (n=6 per sex/genotype). All statistical analyses utilized a two-way linear mixed effects model to look at the effect of genotype within sex and the main effect of genotype across sex. All bars show mean ± SEM. Representative images of the anterior synovium in **(E)** contralateral and **(F)** 7d ACLR limbs. Representative images of the articular surfaces in **(G)** contralateral and **(H)** 7d ACLR joints.

Blinded histopathological scoring was performed to evaluate the effect of TSP2 deficiency on joint structural development and post-injury remodeling at 7d post-ACLR. KO females exhibited marginally lower synovitis scores relative to WT, but no genotype difference was observed in the 7d ACLR limb (Fig.-3C-F**, Suppl.-Fig.-5A**). PTOA composite scores of the entire tibiofemoral joint were significantly higher in the 7d ACLR joints of KO mice, with comparable scores in the contralateral (**Fig.-3C-D**). The higher PTOA score in KO was driven by greater chondrocyte hypertrophy, tibial proteoglycan loss, and femoral subchondral bone thickening (**Fig.-3G-H, Suppl.-Fig.-5B**). These findings show that TSP2-KO mice have greater injury-induced cathepsin activity, indicative of broad inflammation, and accelerated PTOA development at an early post-injury stage.

### TSP2-KO mice exhibit aberrant chronic knee joint remodeling

We next sought to assess how TSP2 deficiency would affect the long-term development of PTOA. Assessing periarticular bone remodeling in uninjured and injured joints, sPLSDA-based dimensionality reduction of all micro-computed tomography (µCT) bone morphometry parameters (**Suppl.-Fig.-6**) identified 3 key components driving sample variation: (1)-sex (45% of variation), (2)-injury (15.2% of variation), and (3)-genotype (11.2% of variation) (Fig.-4A). Genotype-associated variation was driven by increased trabecular spacing and decreased bone volume fraction, trabecular thickness, and bone mineral density in KOs (**Fig.-4B**). These genotype effects persisted when sexes were analyzed separately (**Suppl.-Fig.-7A-D**). Raman spectroscopy in a separate cohort of naïve mice demonstrated reduced collagen crosslinks in trabecular bone, corroborating altered bone matrix in KOs (**Suppl.-Fig.-9A-B**). µCT was further used to assess osteophyte formation, and osteophyte volume was normalized to femoral length given that KO mice had shorter femora (**Fig.-4C**). Unexpectedly, KO males exhibited reduced osteophyte volume compared to WT males, but this was not observed in females (**Fig.-4D-E**).

**Figure 4.**
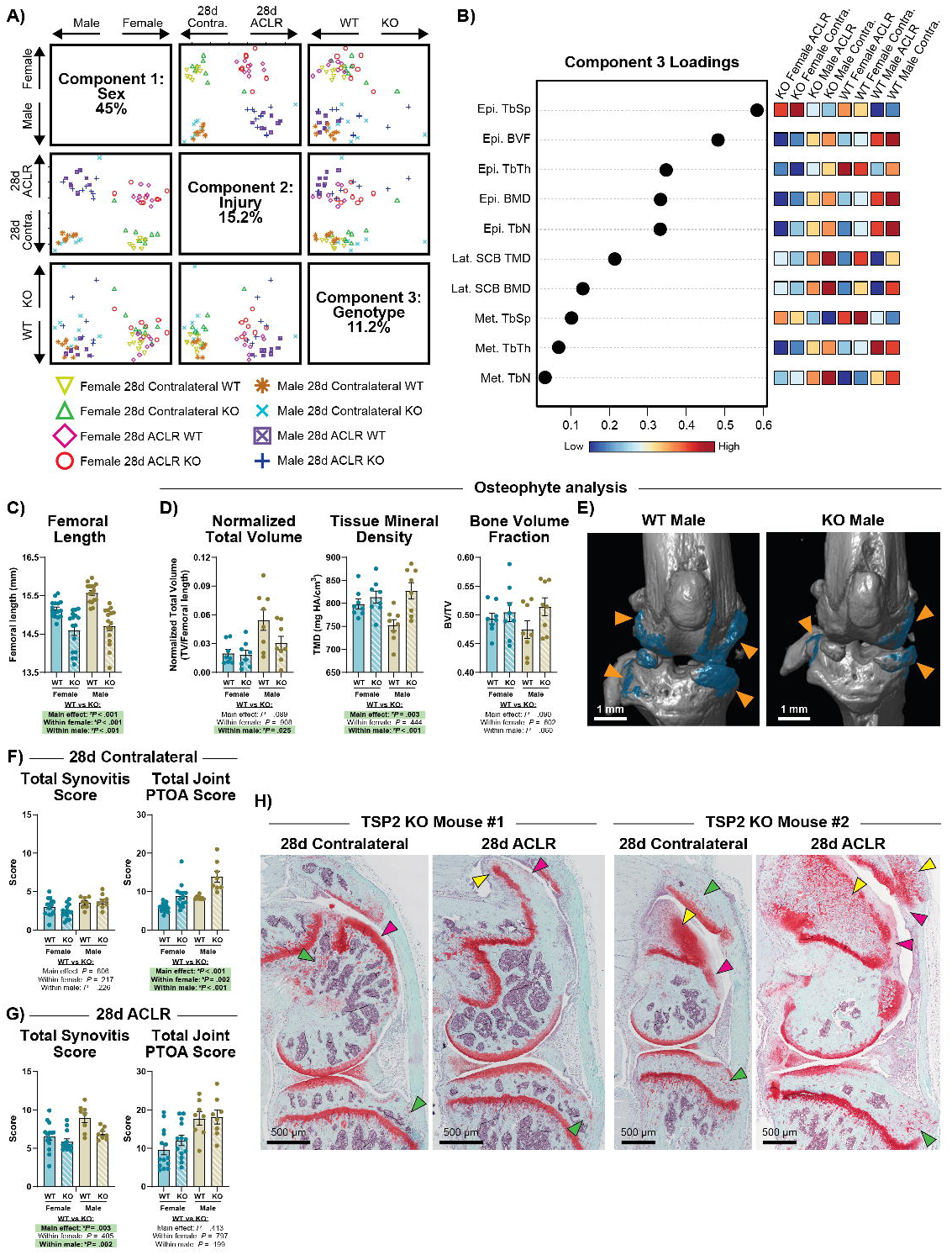
TSP2 KO vs WT 28d post-ACLR bone remodeling, and histopathology. **A)** Overview of sPLSDA analysis of all limbs showing the top 3 components. Each component corresponds to one of the factors in the dataset – sex, injury, and genotype. **B)** Component 3 loadings driving genotype-dependent separation (11.2% of variation). **C)** Femoral length as measured from µCT images. **D)** Osteophyte (OP) analysis of 28d ACLR limbs only includes total OP volume normalized to femoral length, OP tissue mineral density, and OP bone volume fraction (n=8 per sex/genotype). **E)** Representative 3D WT male and KO male samples (blue masks, orange arrows – osteophytes). Histopathological scoring of synovitis severity and PTOA severity in WT and KO **(F)** contralateral and **(G)** 28d ACLR limbs (n=8-14 per sex/genotype). **(H)** Safranin-Orange and Fast Green stained limbs showing aberrant structural remodeling in the contralateral limb and 28d ACLR limb of two mice (green arrow – subchondral bone remodeling, pink arrow – cartilaginous, undermineralized, and acellular surfaces of the patellofemoral region, yellow arrow – large ectopic outgrowths). All statistical analyses utilized a two-way linear mixed effects model to look at the effect of genotype within sex and the main effect of genotype across sex. All bars show mean ± SEM.

Histopathological scoring of the tibiofemoral compartment showed increased PTOA severity in KO contralateral limbs relative to WT contralateral limbs, driven by greater femoral proteoglycan loss, chondrocyte hypertrophy, and, surprisingly, femoral and tibial osteophyte size and maturity (**Fig.-4F, Suppl.-Fig.-8A**). Consistent with our prior studies^40^, contralateral limbs in WT mice did not develop such structural pathology. However, despite these observations in the contralateral joint, no genotype effect on PTOA score was observed in the 28d ACLR limbs (**Fig.-4G**). Synovitis scores were similar in KO contralateral limbs compared to WT contralaterals, but KO 28d ACLR limbs had lower synovitis scores compared to WT 28d ACLRs (**Fig.-4F-G, Suppl.-Fig.-8B**). Broad observation of the entire knee joint, including the patellofemoral region that is not captured in tibiofemoral histopathological scoring, shows extensive aberrant remodeling in TSP2-KO mice in both uninjured (contralateral) and injured (ACLR) states (**Fig.-4H**). Together, these findings demonstrate complex injury responses in peri-articular bone and multiple intra-articular tissues in TSP2-KO mice, suggesting that TSP2 deficiency primes the joint to pathologic structural remodeling that primarily manifests in cartilaginous and bony tissues.

### Mature, naïve TSP2-KO mice exhibit greater idiopathic OA severity

After observing that uninjured contralateral joints of 12-14 wk-old KO mice exhibited degenerative changes not observed in uninjured WT joints, we sought to assess whether mature KO mice would differentially develop idiopathic OA-like changes. Aged female WTs and KOs (36-42 wks old) were assessed for idiopathic alterations in naïve joints. Bone morphometry analysis of µCT images revealed inferior bone quality in KOs, driven by decreased tissue mineral density and trabecular thickness (**Suppl.-Fig.-10A-F**). Histopathologic analysis showed that naïve mature KO mice had similar synovitis but worse OA scores compared to WTs, driven by increased subchondral bone thickness, tibial proteoglycan loss, femoral structural damage and chondrocyte hypertrophy (**Suppl.-Fig.-10G-H**). These findings further demonstrate that TSP2 deficiency accelerates development of an OA-like phenotype.

### Uninjured TSP2-KO mice exhibit a PTOA-like synovial transcriptome

Given the synovial origin of injury-induced TSP2, we last performed bulk RNAseq of synovium in Sham, 7d ACLR, and 28d ACLR WTs and KOs. Samples underwent rigorous quality control, and RNA integrity and sequencing quality control metrics were comparable across final samples included in analyses (**Suppl.-Tables 4-7, Suppl.-Fig.-11A-B**). PCA and Eigencor analyses showed that injury and genotype were the major drivers of sample variance; consequently, sexes were merged for analysis (**Suppl.-Fig.-11A, 11C**). Comparing WT vs KO synovia within each group, we observed the greatest number of DEGs (*P_adj_* < .05) in the Sham WT vs KO comparison (2,089), with far fewer in 7d ACLR (119) and 28d ACLR (20) groups (Fig.-5A-B**, Suppl.-Fig.-12A-B**). Gene set enrichment analysis compared Sham KO to Sham WT (**Fig.-5C**) and demonstrated upregulation of terms indicative of immune activation (“leukocyte-mediated immunity”, “cytokine-mediated signaling pathway”), oxidative stress (superoxide and reactive oxygen species metabolic processes), and cytoskeleton remodeling (“actin filament organization”) alongside downregulation of mitochondrial metabolism-related pathways, indicative of metabolic reprogramming (“proton motive force-driven ATP synthesis”, “tricarboxylic acid cycle”). “Angiogenesis” was also increased in KO Sham relative to WT, consistent with TSP2’s anti-angiogenic role (**Fig.-5C**). Orthogonal Reactome pathway analysis extended these findings showing upregulation of VEGF signaling, as well as terms related to neuronal signaling and nerve sprouting (“nervous system development”, “axon guidance”), particularly in KO Shams (**Suppl.-Fig.-12C**). Within the 7d ACLR group, KOs were enriched in pathways related to lipid metabolism (“fatty acid metabolic process”, “lipid oxidation”, “peroxisome organization”), suggesting active remodeling and inflammatory activation of the synovial adipose tissue. 28 ACLR KO synovium showed increased stress response (“ER-nucleus signaling pathway”, “ubiquitin-dependent ERAD signaling pathway”) compared to WT (**Fig.-5C**).

**Figure 5.**
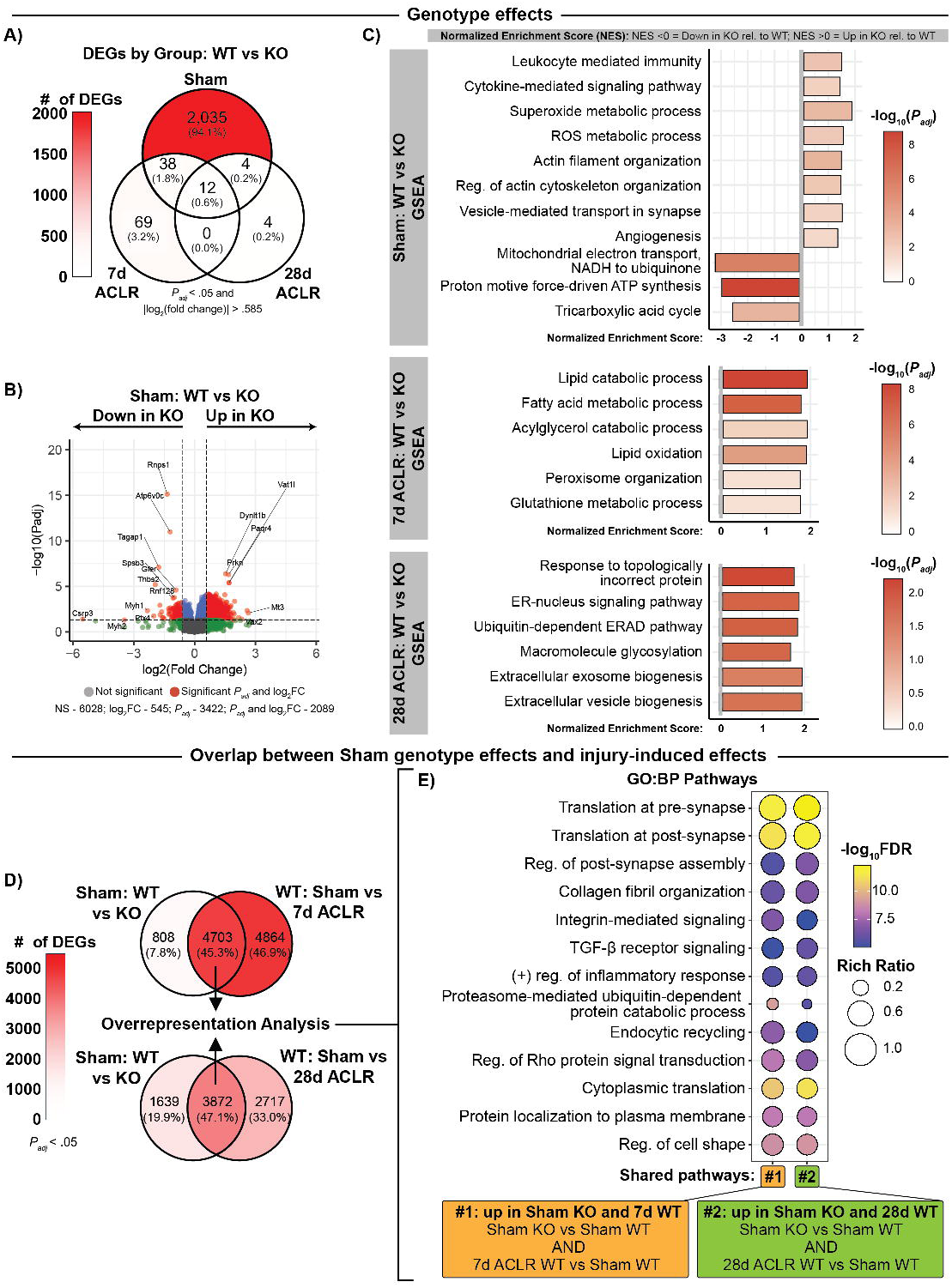
Bulk RNAseq of TSP2 KO and WT synovium. Global TSP2 knockdown results in a PTOA-like phenotype in the synovial transcriptome. **A)** Three-way venn diagram showing DEGs between WT and KO synovium across the three different groups – Sham, 7d ACLR, and 28d ACLR (*P_adj_* > .05, |log2FC| > .585). **B)** Volcano plot showing the highest number of DEGs when comparing WT and KO synovia within the Sham group. **C)** GSEA of WT vs KO synovia within each group with a curated list of terms shown. **D)** Venn diagrams showing overlapping DEGs between the Sham genotype effect and the early-stage WT injury effect or late-stage WT injury effect (*P_adj_* > .05). **E)** PantherDB gene ontology analysis of shared DEGs between Sham KO vs Sham WT and WT 7d ACLR vs WT Sham (column #1) and Sham KO vs Sham WT and WT 28d ACLR vs WT Sham (column #2). A curated list of terms shown is shown. N=2-3 per sex/group/genotype.

Given the aberrant structural remodeling we observed in uninjured KO joints, we evaluated overlapping DEGs between the Sham genotype comparison, which isolates the baseline effect of TSP2 deficiency, and injury-induced genes in WT mice. There was substantial overlap in the DEGs of these two comparisons, with the Sham genotype effect sharing 45-47% of its DEGs with the WT injury effect (**Fig.-5D**). Gene ontology analysis revealed a strong shared synapse-related (translation at the pre- and post-synapse, “regulation of post-synapse assembly”), ECM and fibrosis-related (“collagen fibril organization”, “integrin-mediated signaling”, “TGF-β receptor signaling”), and inflammatory response profile between the Sham genotype effect and the WT injury-induced effect (**Fig.-5E**). Pathways overlapping uniquely with the 7d ACLR effect include tissue activation and remodeling pathways (“bone development”, “regulation of fibroblast proliferation”, “regulation of stem cell differentiation”). Pathways overlapping uniquely with the 28d ACLR effect include “angiogenesis” and mechanotransduction-related pathways (**Suppl.-Fig.-12D**). These findings suggest that TSP2 deficiency primes the synovium towards a pathological phenotype, driving susceptibility to accelerated OA and aberrant joint remodeling.

## DISCUSSION

Here we demonstrate that TSP2, a matricellular protein with diverse anti-angiogenic and ECM assembly functions, is strongly induced by sublining synovial fibroblasts following joint injury and that genetic TSP2 deficiency results in aberrant injury responses and accelerated age-related joint degeneration.

We observed rapid and sustained induction of TSP2 in sublining fibroblasts after injury, and our *in vitro* results suggest that this may be driven, at least in part, by overactive TGF-β and Wnt signaling activity, which we have also shown to be activated by injury^40,41^. Prior work in umbilical cord blood-derived MSCs indicates TSP2 negatively regulates Wnt/β-catenin and TGF-β signaling^49^. Combined with our finding that WNT3a and TGF-β upregulate *Thbs2* in synovial fibroblasts, this suggests TSP2 may mediate a negative feedback loop for Wnt signaling and TGF-β signaling. Our RNAseq data supports increased TGF-β activity in TSP2-KO synovia, but further studies are necessary to assess whether the pathological manifestations in TSP2-KO joints are due to overactivity of these pathways previously linked to OA and PTOA pathogenesis.

Our findings strongly support that TSP2 is necessary for normal joint homeostasis and aging, as TSP2-KOs exhibited aberrant remodeling in uninjured joints. RNAseq of the synovium corroborates these observations, as a major finding of our study was an injury-like transcriptome in uninjured TSP2-KO synovium, suggesting that KO mice are primed for injury responses. A direct comparison between the uninjured KO synovium and injured WT synovium showed substantial overlap in genes and associated disease-relevant processes including inflammatory and innate immune activity, fibrosis-relevant terms including collagen production and TGF-β signaling, and synapse-related processes. Direct comparisons of uninjured TSP2-KO and WT synovium revealed enriched angiogenesis and inflammatory processes, with correspondingly increased ROS activity and synaptic vesicle trafficking. Furthermore, we observed strongly downregulated terms related to oxidative phosphorylation in TSP2-KO synovium, suggestive of dysfunctional mitochondrial metabolism. As TSP-2 is a secreted matricellular protein with no described intracellular role, further studies should assess how its deficiency results in the injury signature and metabolic reprogramming that may underpin the aberrant structural remodeling in TSP2-KO joints.

TSP2 mice exhibited worse histopathological PTOA severity at 7d post-ACLR, indicating more rapid onset of PTOA, which is consistent with TSP2-KO mice exhibiting a disease-primed joint. By 28d post-ACLR, PTOA scores were similar across genotypes; however, despite this, OP size was lower in male KOs versus WTs. A role for TSP2 in endochondral ossification, the process by which OPs form, is supported by fracture studies demonstrating that TSP2-KO mice exhibit accelerated callus bone formation, driven by diminished endochondral ossification and increased appositional bone formation^50^. Our observation of lower OP volume in TSP2-KO mice is consistent with these prior findings and supports TSP2-dependent regulation of endochondral ossification in OP formation. While our RNAseq suggests overactive angiogenesis-related pathways in KOs, Angiosense imaging was similar across genotypes. Thus, we cannot conclude whether the effects we observed in TSP2-KO are due to modulation of tissue vascularity. Limited evidence in a rheumatoid arthritis-focused study showed that TSP2 overexpression decreased vascularization of implanted human synovium in SCID mice^20^. Furthermore, the overactive immune and inflammatory processes we observed in uninjured TSP2-KO joints may also drive greater disease severity after injury, which partially persisted up to 28d post-ACLR. Considering the massive KO effect in uninjured limbs, future studies using inducible or cell-specific TSP2-KO models are needed to isolate TSP2’s role in injury-induced pathology. Future studies should also explore the potential for TSP2 mimetics as therapeutics for PTOA.

In summary, our findings demonstrate that the matricellular protein TSP2, found to be highly expressed by synovial fibroblasts after joint injury, plays important roles in joint homeostasis and injury responses.

TM is a consultant for Relation Rx

## Funding

National Science Foundation Graduate Research Fellowship grant DGE 2241144 (LL)

Congressionally Directed Medical Research Programs (HT9425-23-1-0327)

National Institutes of Health-National Institute of Arthritis and Musculoskeletal and Skin Diseases (NIH NIAMS) (1R21AR076487-01A1)

AJK was supported by a Pioneer Fellowship from the University of Michigan, and a K99 Award from the National Institutes of Health - National Institute of Arthritis and Musculoskeletal and Skin Diseases (NIH NIAMS) (K99AR081894)

## Acknowledgements

The authors would like to acknowledge the Michigan Integrative Musculoskeletal Health Core Center (MiMHC) which is supported by a P30 grant (P30 AR069620) from NIH/NIAMS for processing samples for histology and for use of their microscopy equipment. We also wish to acknowledge the University of Michigan Biomedical Research Core Facilities, specifically the flow cytometry core, which is a part of the Medical School Office of Research, for use of their flow cytometer.

## Author contributions

Conceptualization: LL, AIA, KDH, TM

Methodology: LL, GM, TM

Investigation: LL, LMJ, AJK, MDN, HD, CRD, AM, IJS, SCH, GM, SC, SGN, SH

Visualization: LL, AJK, HD, AM, IJS, SCH

Funding acquisition: LL, AJK, AIA, KDH, TM

Writing – original draft: LL, TM

Writing – review & editing: LL, AJK, MDN, AIA, KDH, TM

## Data and materials availability

Bulk RNA-seq data will be publicly available on GEO and all other data are available in the main text or the supplementary materials.

---

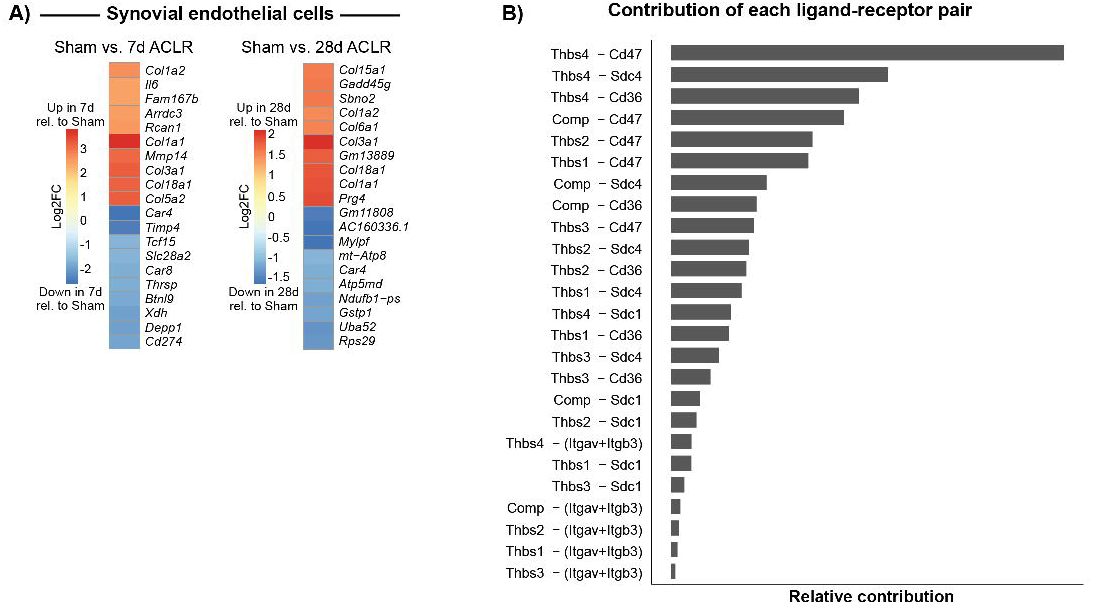

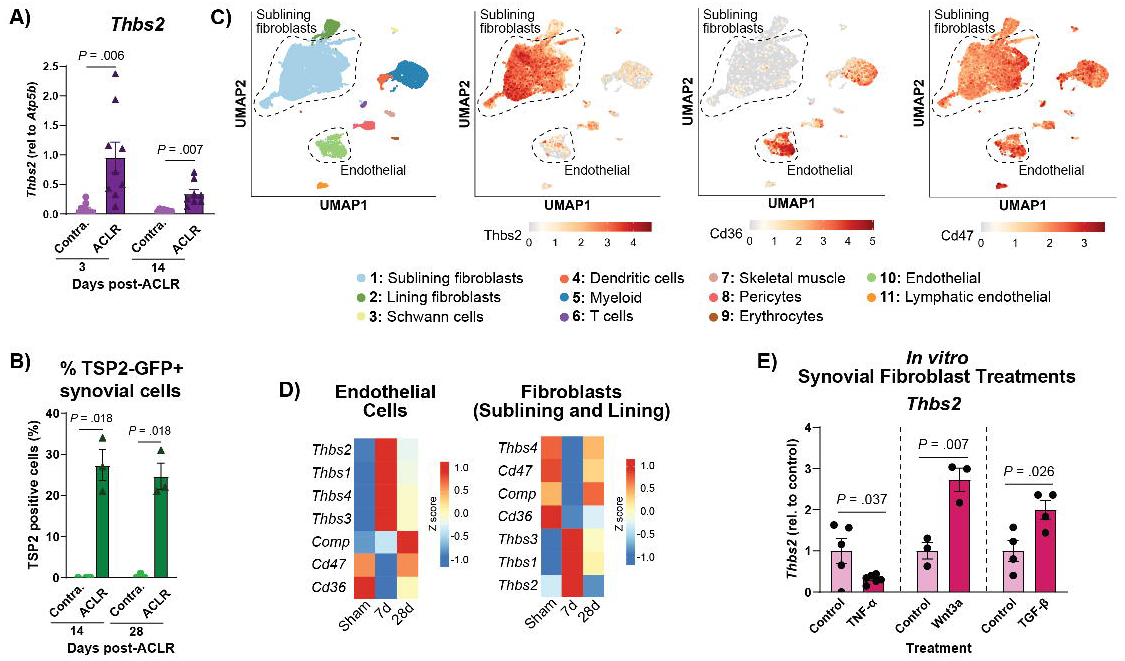

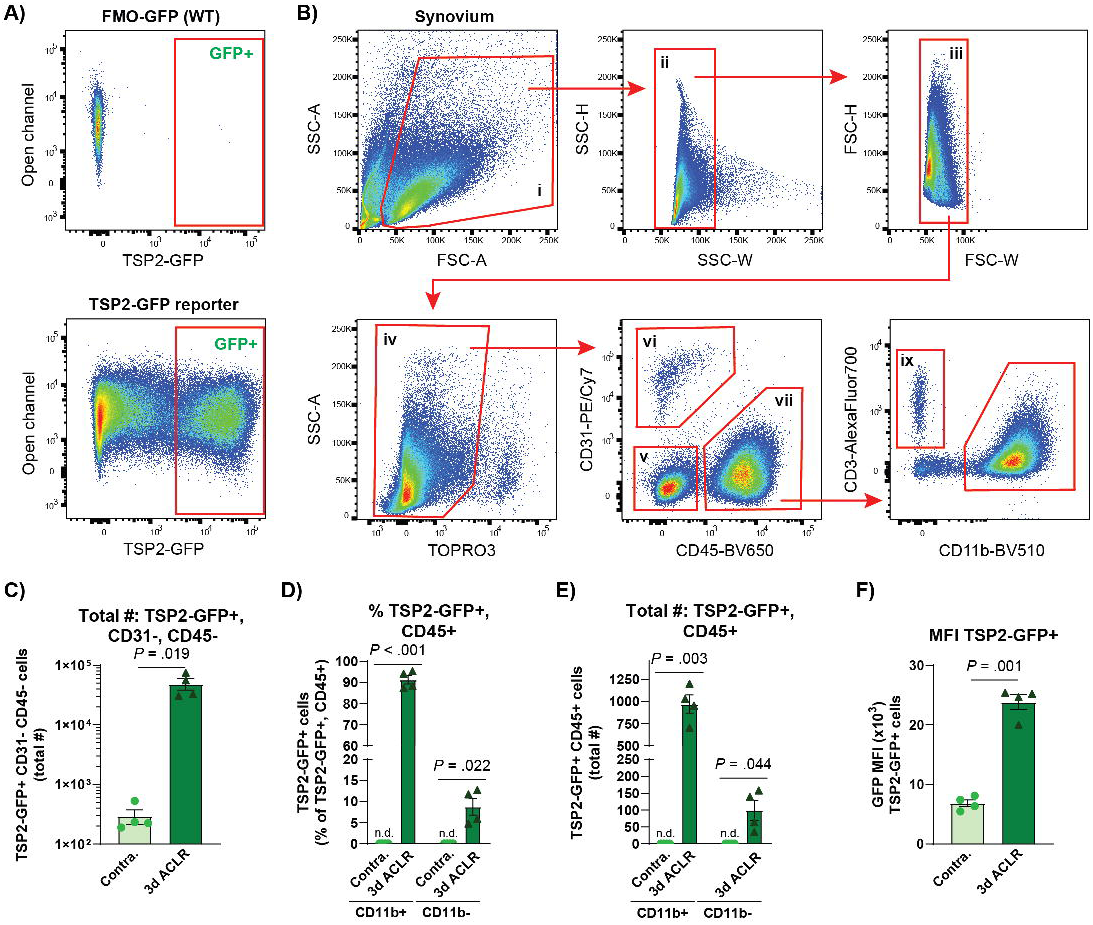

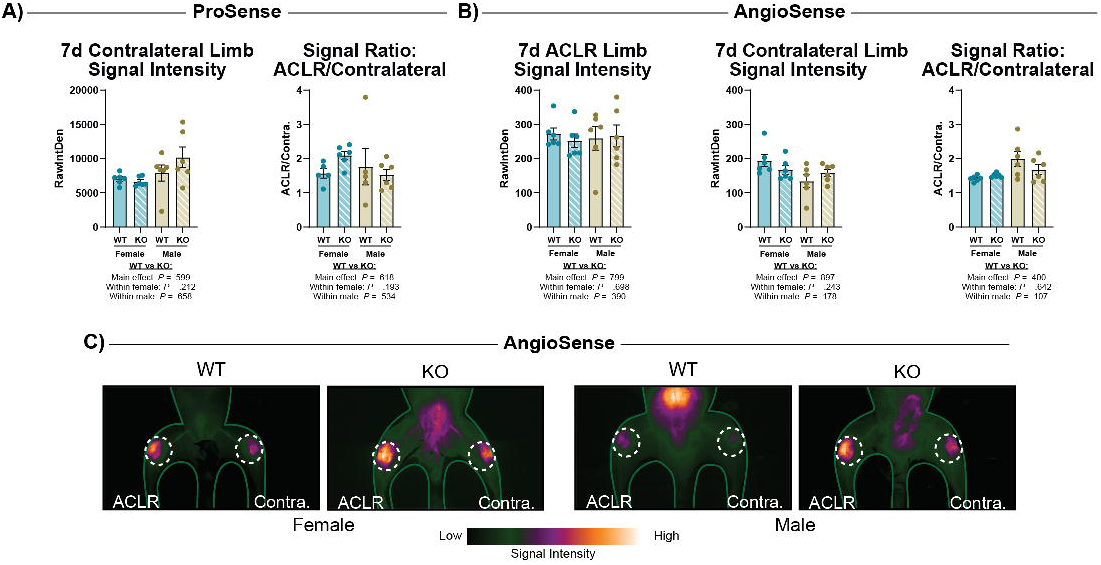

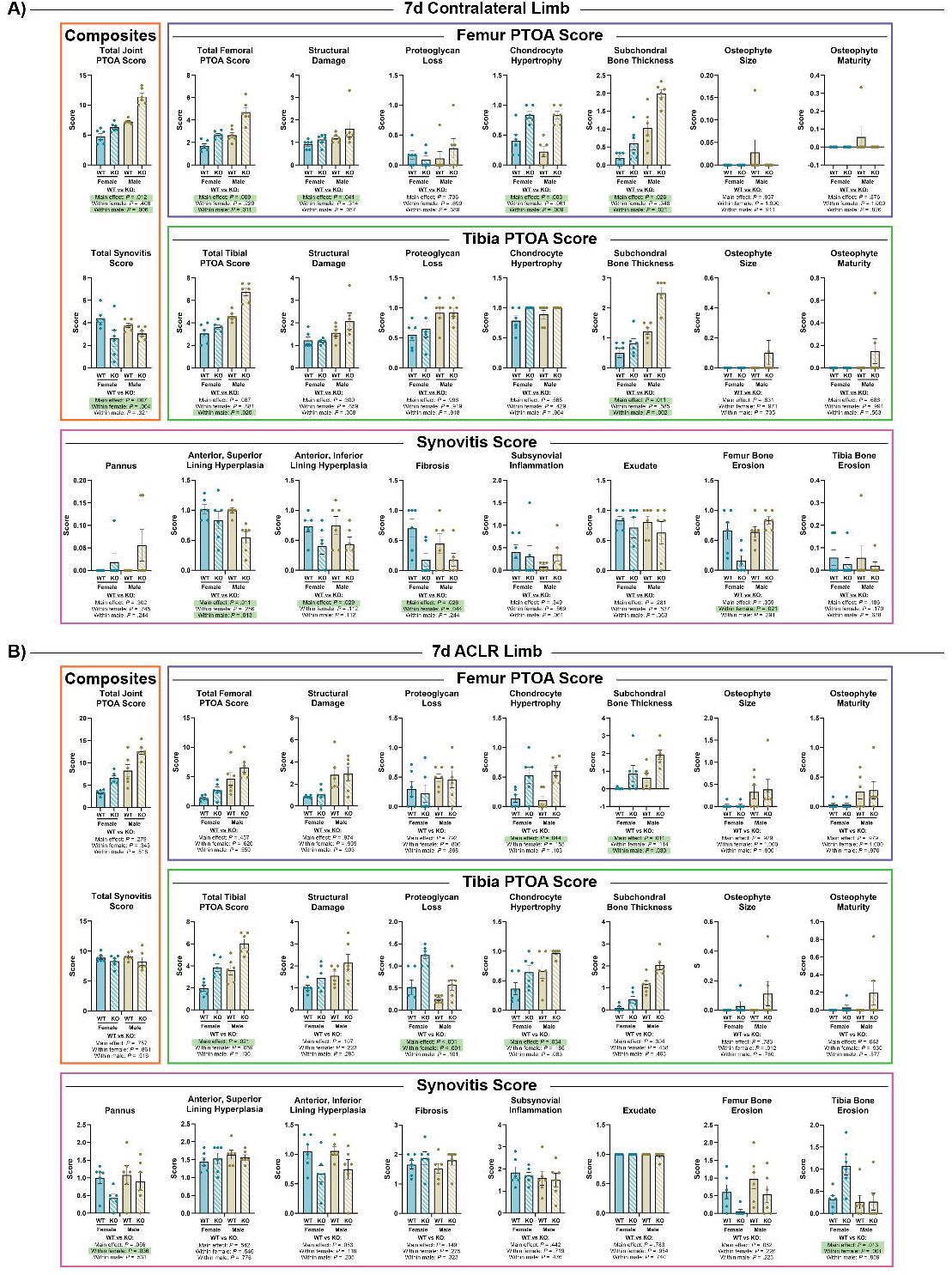

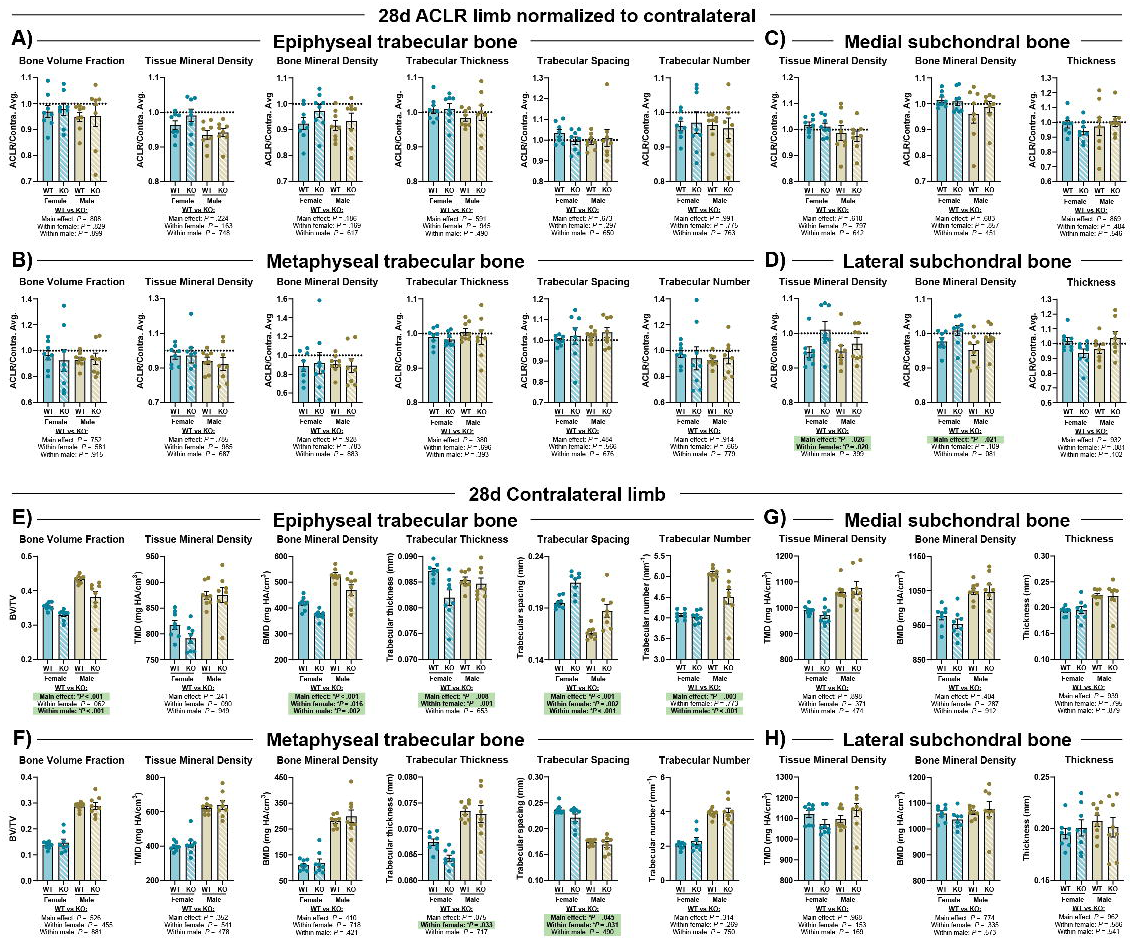

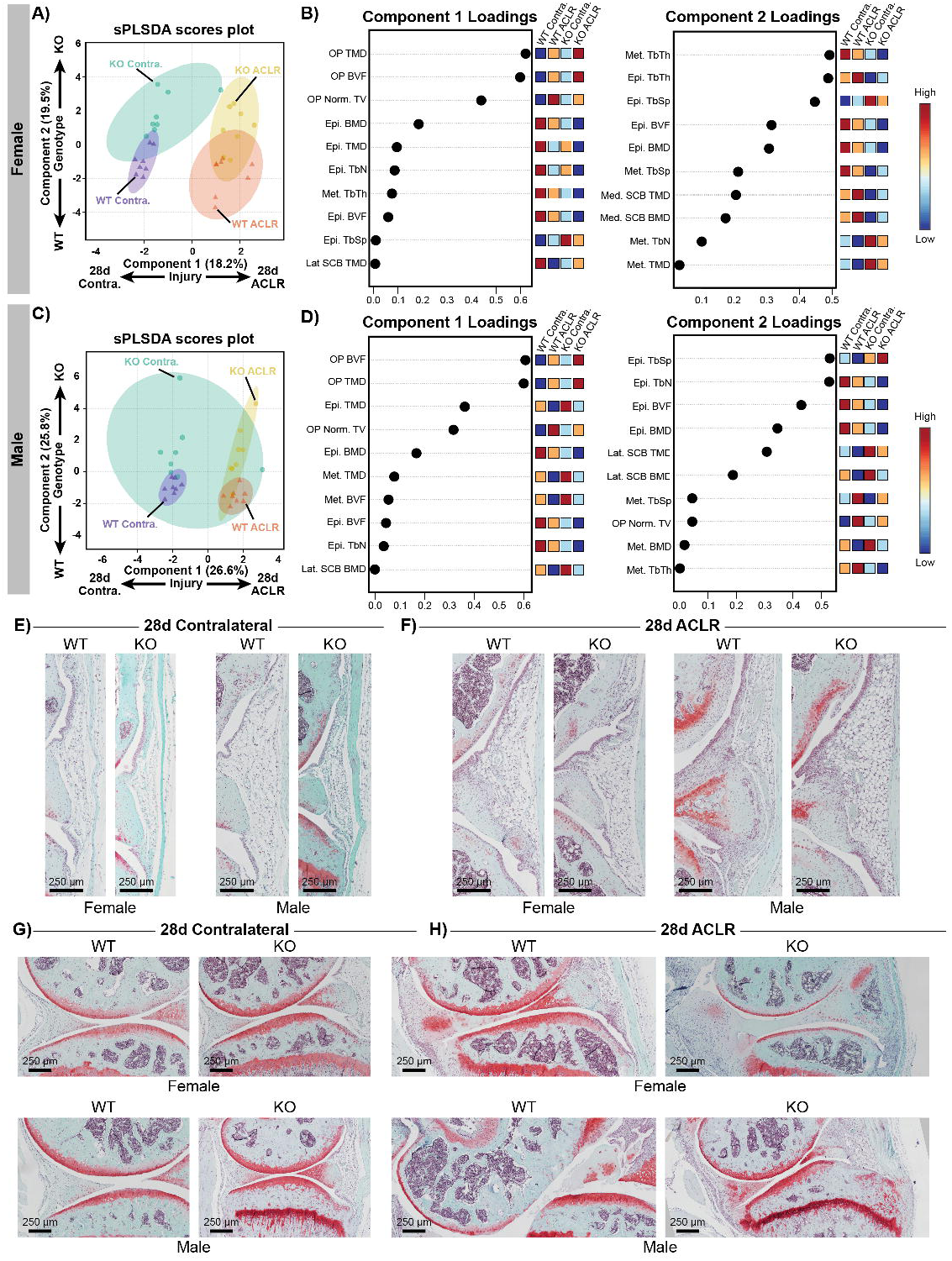

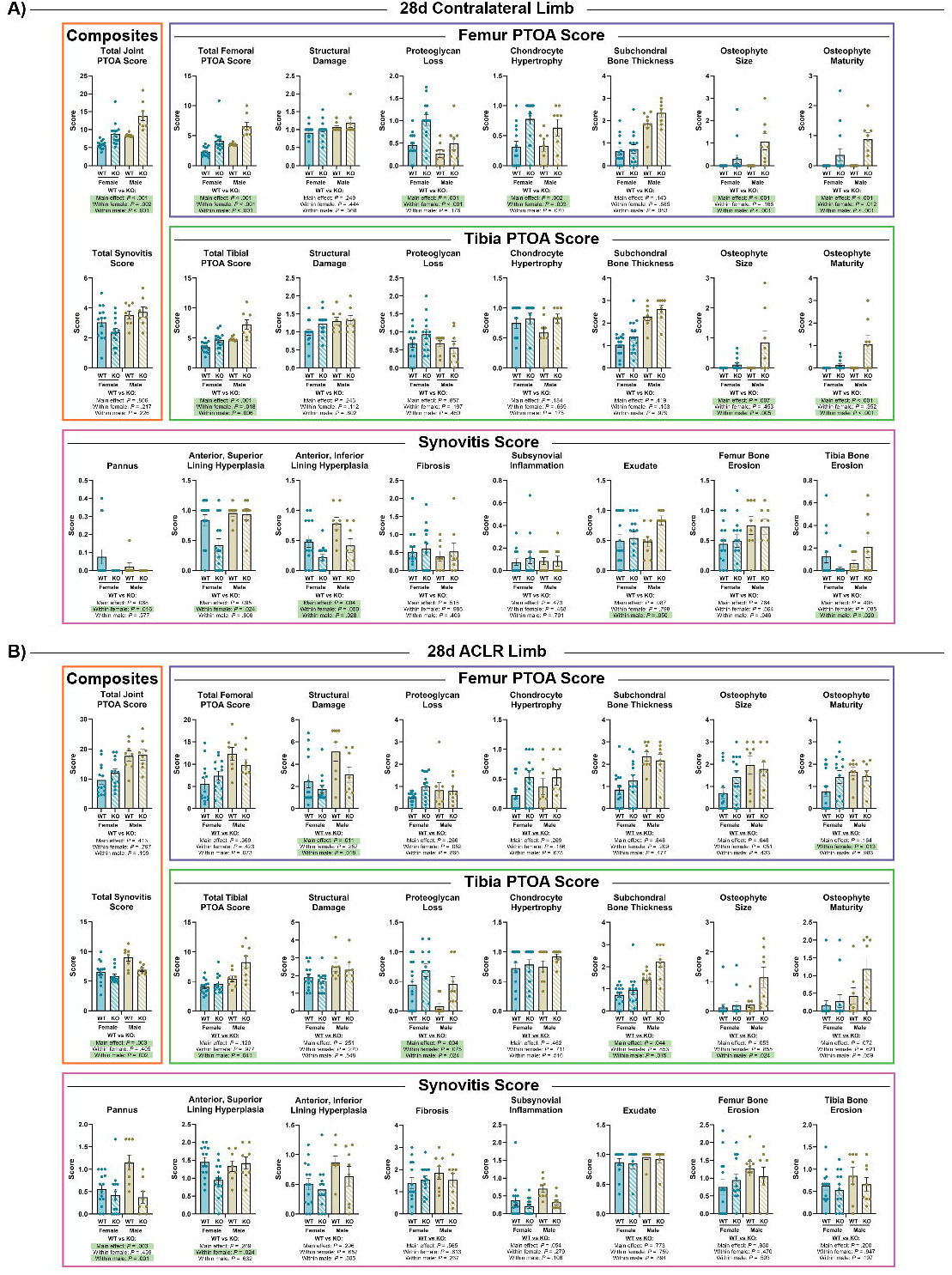

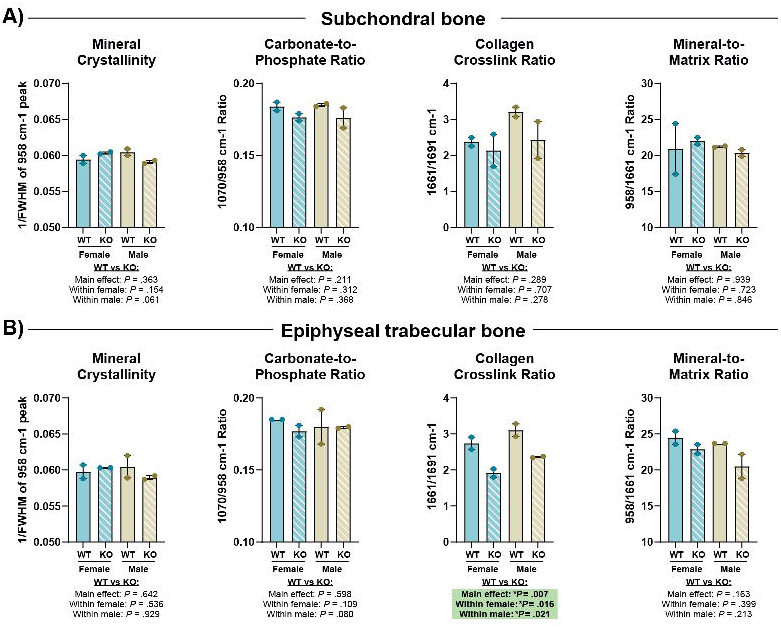

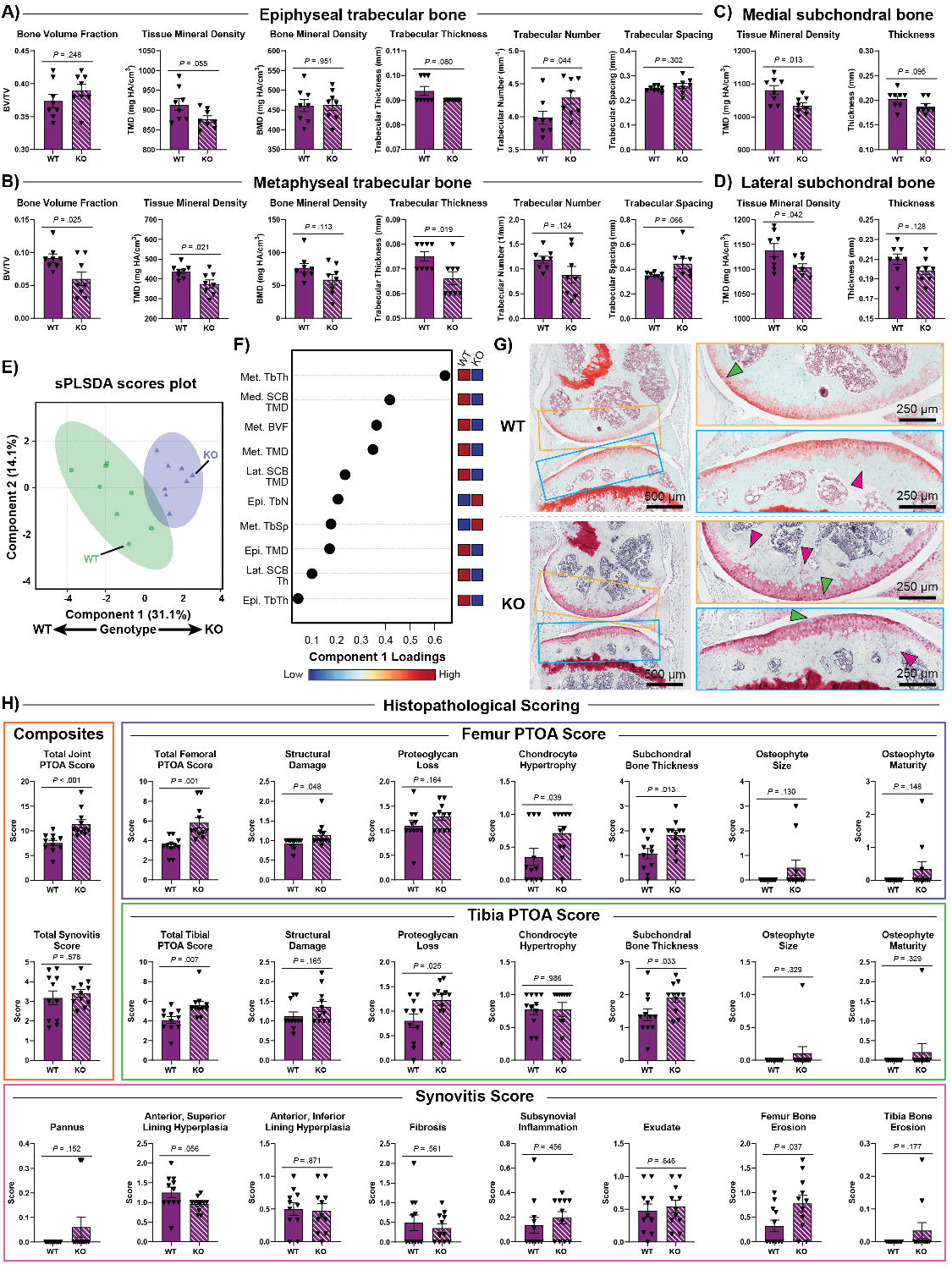

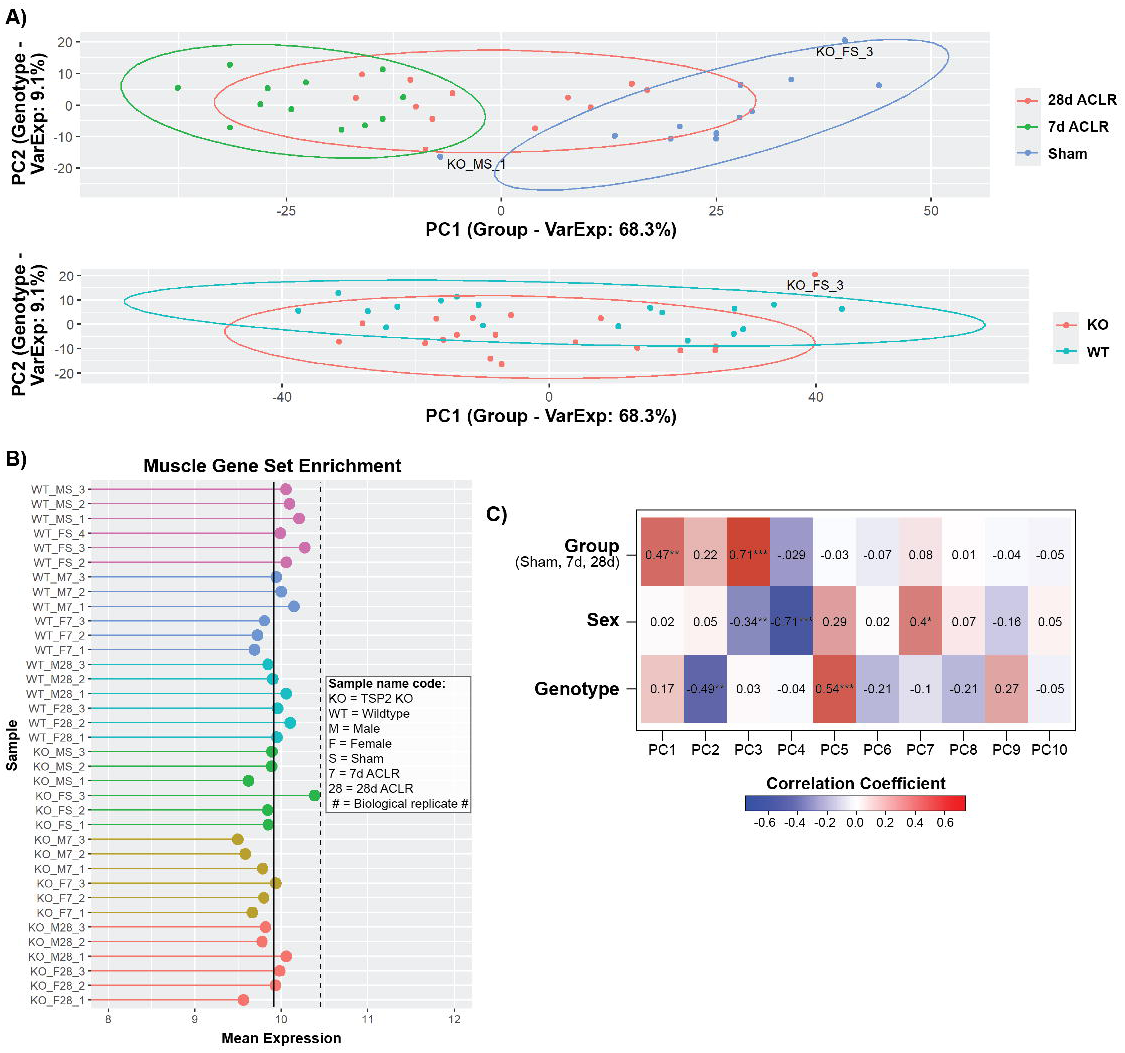

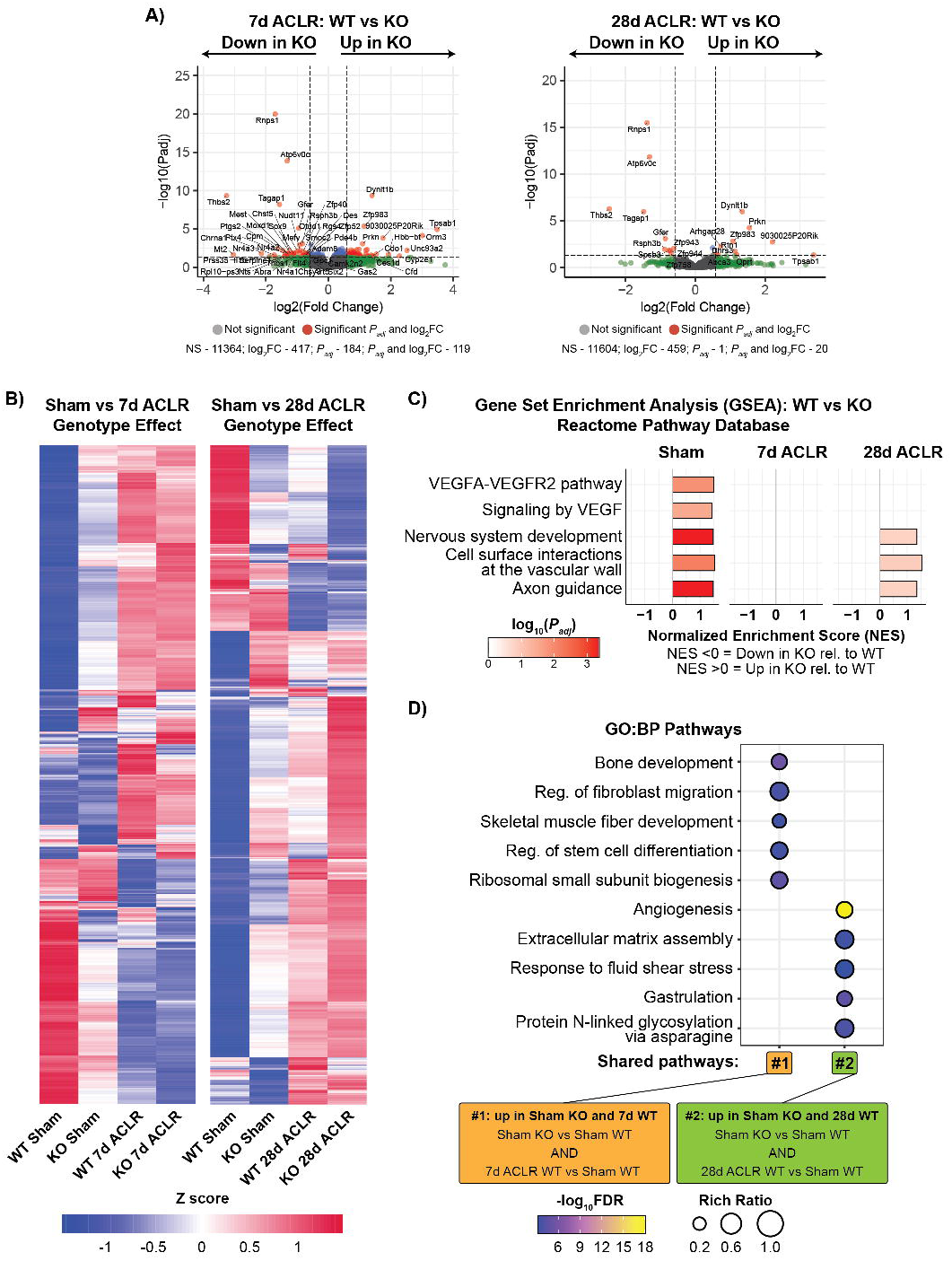

