## Supplemental Materials and Methods for "Thrombospondin-2 deficiency primes the synovial joint for aberrant tissue remodeling and injury response"

*Mice and PTOA model*

Non-invasive anterior cruciate ligament rupture (ACLR) was utilized to induce PTOA as described previously[1]. Briefly, under isoflurane anesthesia (5% induction, 2-2.5% maintenance), mice were positioned in a prone position on a mechanical testing system (ElectroForce 3300AT, TA Instruments, New Castle, DE, USA) situated with a custom fixture to hold the right knee at 100° of flexion. Following a preconditioning loading cycle, a rapid 1.5 mm displacement caused an isolated, mid-substance rupture of the anterior cruciate ligament. Sham animals underwent a cycle of anesthesia and analgesia without any mechanical loading. For all injury studies, mice were randomized to Sham or injury group.

*Near-infrared (NIR) imaging*

<u>AngioSense 750 in WT mice 7d and 28d post-ACLR:</u> At 7d or 28d post-ACLR, WT mice were anesthetized and injected retro-orbitally (31-gauge needle, BD SafetyGlide Insulin Syringe) with AngioSense 750 (100 μL per mouse).

<u>ProSense 680 + AngioSense 750 in WT and KO mice 7d post-ACLR:</u> A separate cohort of WT and KO mice underwent ACLR and at 7d post-ACLR received intra-articular ProSense 680, also known as Pan Cathepsin 680, injections (4 μL per limb) and retro-orbital AngioSense 750 injections (100 μL per mouse).

<u>NIR imaging and analysis:</u> Four hours following injection, mice were euthanized, skinned, and near infrared imaging was conducted at 700nm and 800nm wavelengths as appropriate (Pearl Impulse, LI-COR, Lincoln, Nebraska). Using Fiji – ImageJ, consistently sized regions of interest were drawn over each knee, and the raw integrated density (RID) was

recorded. Both RID and the RID ratio which normalized ACLR RID/Contralateral RID are reported.

*Mining of publicly available transcriptomics datasets*

Using our published mouse synovial scRNAseq atlas[2], we identified the top ten upregulated and downregulated genes in endothelial cells comparing the early injury (7d ACLR) and late-stage injury (28d ACLR) effects relative to Sham. This data was also mined to examine how all synovial cells and more specifically fibroblasts and endothelial cells change their expression profile of the thrombospondin family of genes (*Thbs1, Thbs2, Thbs3, Thbs4, Comp)* and the TSP2 receptors, *Cd36* and *Cd47* following ACLR.

The gene set for the "Angiogenesis" GO term was downloaded from the AmiGO 2 database. Using the "Angiogenesis" GO term, we looked at overlapping genes between the "Angiogenesis" gene set and differentially expressed genes (DEGs, $P_{adj} < 0.05$) from our previously published bulk RNA seq dataset comparing Sham vs 7d ACLR or Sham vs 28d post-ACLR mouse synovium[1]. We then looked at the expression of these overlapping genes in our published mouse synovial scRNAseq atlas[2], to identify which synovial cell types were responsible for driving the DEGs in the "Angiogenesis" GO term. The bulk RNAseq data was used for the initial identification of "Angiogenesis" DEGs as bulk RNAseq has greater sequencing depth as compared to scRNAseq. The Z-score of expression of the overlapping genes in scRNAseq were plotted by synovial cell type using the aggregate expression of genes.

Using the same mouse synovium scRNAseq dataset[2], we removed erythrocyte and skeletal muscle clusters and then performed cellular communication analyses using CellChat[3], which infers cellular crosstalk based on the expression of established ligand-receptor pairs.

Outgoing communication patterns were calculated (k=8) and visualized using a river plot with each cellular subset coordinating a distinct signaling module, termed a "pattern". Within the pattern mediated by synovial sublining fibroblasts was the thrombospondin (THBS) signaling pathway network. A heatmap of the THBS signaling pathway network was visualized using the *netAnalysis_signalingRole_network()* function, allowing visualization of which synovial cells were responsible for influencing, mediating, receiving, and sending thrombospondin signals. The contribution of all ligand-receptor pairs in the THBS signaling pathway, which includes the ligands *Thbs1*, *Thbs2*, *Thbs3*, *Thbs4*, and *Comp*, along with their receptors, was visualized using *netAnalysis_contribution()*. The primary *Thbs2* ligand-receptor pair signaling networks (*Thbs2*-*Cd36*, *Thbs2*-*Cd47*) were visualized using circle plots to evaluate the strength of intra- and intercellular *Thbs2* signaling in a cell specific manner.

Our scRNAseq data[2] was again utilized to identify *Thbs2*, *Cd36* and *Cd47* expressing cells. Feature plots showing the expression of *Thbs2, Cd36,* and *Cd47* across the all cells, all conditions merged object, which contains integrated Sham, 7d ACLR, and 28d ACLR synovial cells were generated using the *Shiny* R package. Endothelial cells and fibroblasts (sublining and lining together) were separately subset from all synovial cells across all conditions to examine expression of the thrombospondin family genes and the receptors *Cd36* and *Cd47.*

*Flow cytometry*

TSP2-expressing cells were analyzed using TSP2-eGFP reporter mice, generated previously[4]. Briefly, synovia were micro-dissected from the anterior, lateral, and medial knee joint compartments including infrapatellar fat pad, and enzymatically and mechanically digested to yield single-cell suspensions as previously described[2,5], using 400 μg/mL final concentrations of Liberase TM, Collagenase IV, and DNaseI, in DMEM media. Contralateral and injured joints were harvested separately, and each sample consisted of a single synovium (n=4 ACLR, n=4 contralateral, equal sex representation). Tissue from C57BL/6J mice were used as WT controls, lacking endogenous GFP.

After digestion, cells were pre-blocked with Mouse TruStain FcX PLUS (1:1000, Biolegend Cat. #156604) in FACS buffer (PBS containing 2% FCS and 1 mM EDTA). Cells were then incubated for 30 min in a dark cold room with a cocktail of fluorescently conjugated antibodies, available in **Suppl.-Table 1**. Unstained cells, single-stained cells, and fluorescence-minus-one (FMO) cells were included as experimental controls. TOPRO3 dye (1:100,000) was added to cells prior to running flow cytometry on a BD LSR Fortessa machine. FlowJo v10 (BD/Treestar) was used for analysis.

**Suppl. Table 1. Flow cytometry antibodies.**

| Antibody | Clone (RRID) | Vendor | Application | Dilution |
|---|---|---|---|---|
| Anti-mouse CD3 AlexaFluor700 | 500A2 (AB_396972) | BD Pharmingen | Flow cytometry | 1:100 |
| Anti-mouse CD11b Brilliant Violet 510 | M1/70 (AB_2561390) | Biolegend | Flow cytometry | 1:400 |
| Anti-mouse CD31 PE/Cy7 | 390 (AB_830756) | Biolegend | Flow cytometry | 1:200 |
| Anti-mouse CD45 Brilliant Violet 650 | 30-F11 (AB_2565884) | Biolegend | Flow cytometry | 1:400 |

*Cryohistology*

Hindlimbs form TSP2-GFP mice were removed via disarticulation at the femoroacetabular joint. TSP2-GFP reporter limbs were fixed in 10% NBF for 48 hours while nutating at 4ºC. Decalcification was performed in 10% EDTA for 2 weeks while nutating at 4ºC. Sucrose cryo-preservation was then performed and samples were embedded in Optimal Cutting Temperature (OCT) compound. Joint tissue sections (10 µm thick) of the medial condyle were cut in the sagittal plane. Sections were stained with DAPI nuclear dye and imaged (10X magnification, Agilent, BioTek Lionheart FX, Santa Clara, CA) to observe distribution of TSP2 throughout the joint.

*Synovial tissue gene expression via RT-qPCR*

Upon thawing, samples were homogenized in Trizol using an orbital homogenizer (Precellys Evolution, Bertin Instruments, Rockville, MD, USA) and CK28 Precellys homogenization tubes. RNA was isolated via chloroform phase separation and converted to cDNA (High Capacity cDNA Reverse Transcription Kit, Applied Biosystems, CA, USA). cDNA was combined with SYBR Green master mix (Applied Biosystems, MA, USA) and relevant *Thbs2* gene primers for RT-PCR with a Bio-Rad T100 Thermal Cycler (Bio-Rad, Berkeley, CA, USA). Data was normalized to the housekeeping gene, *Atp5b*, using the $2^{-\Delta CT}$ method.

*Xenium Prime 5K spatial transcriptomics*

Sample collection and processing was performed as extensively detailed in our previously published protocol[6]. In brief, under deep isoflurane anesthesia (5% induction, 3% maintenance), transcardiac perfusion was performed with cold PBS (20 mL) followed by Z-fix zinc formalin fixative (50 mL) at a rate of 200 mL per hour. Hindlimbs were removed and fixed for an additional 48 hrs in Z-fix while nutating at 4ºC. Samples were rinsed under running tap water then decalcified in 80% Immunocal in PBS for 16 hrs while nutating at 4ºC. Samples were again rinsed and submitted to the Michigan Integrative Musculoskeletal Health Center's Structure, Composition, and Histology Core for paraffin processing.

Each sample was individually paraffin embedded, trimmed, and oriented in the sagittal plane. Tissue sections were collected to extract RNA using a Qiagen RNeasy FFPE isolation kit and the RNA quality was assessed using an Agilent Bioanalyzer. Individual sample paraffin blocks were melted down and samples were trimmed such that six joints could then be co-embedded into a single block that was the size of the Xenium slide capture area. In total, two co-embedded blocks were generated, each with one joint per sex (male, female) per group (Sham, 7d ACLR, 28d ACLR). One section from each block was collected onto a Xenium slide and submitted for spatial transcriptomics processing. All sectioning took place in a dead airbox that was UV-sterilized and cleaned with RNase Zap.

As published previously[6], spatial transcriptomics processing used the 10X Genomics Xenium Prime 5K Mouse Pan Tissue and Pathways Panel, a custom 100-gene musculoskeletal tissue-focused panel, and the optional multimodal cell segmentation cocktail. Hematoxylin and eosin (H & E) staining was performed after the Xenium Analyzer run. The H & E image was imported into Xenium Explorer and aligned to the Xenium morphology image that was generated from the multimodal cell segmentation. The anterior and posterior synovia were manually outlined and the cells within the region of interest (ROI) were computationally subset in Xenium Explorer. High quality transcripts (Q-value $\geq$ 20) within the ROIs were used for further analyses. All synovial outlines were integrated using SCTransform normalization and RPCA integration in Seurat. Broad cell types and fibroblast subsets were identified using

*FindAllMarkers()*. Cell IDs and their annotations were exported from R and imported into Xenium Explorer for visualization. Sham and 28d ACLR limbs from male mice were chosen to generate visualizations with endothelial cells and fibroblasts annotated along, with transcript overlays of *Thbs2*, *Cd36* and *Cd47*.

*Micro-computed tomography (μCT)*

Contralateral and ACLR limbs from WT and KO mice were harvested at 28d post-ACLR (n=8 per sex/genotype) and a separate cohort of naïve 36-42 wk old (n=8 genotype) and WT were harvested to evaluate the effect of TSP2 deficiency on injury-induced periarticular bony remodeling and age-related changes in bone structure. Prior to scanning, fixed limbs were rehydrated in PBS, imaged (SkyScan 1176; 8.9-μm voxel), and reconstructed. Femoral metaphyseal, epiphyseal, and medial and lateral subchondral bone compartments were automatically contoured for bone morphometry analysis using a Matlab-based neural net segmentation algorithm and manually checked and edited. Femoral length was measured along the longest axis of the femur, and osteophytes were manually contoured and analyzed using Dragonfly.

Sparse partial least squares discriminant analysis (sPLSDA) in MetaboAnalyst was utilized for dimensionality reduction of raw microcomputed tomography output parameters. For the 28d ACLR cohort, sPLSDA (5 components, 10 variables per component) was performed on all limbs (contralateral/ACLR, WT/KO, female/male) and an overview of the top 3 components separating the data was output. The top 10 loadings were shown for Component 3 which was driven by genotype effects. As Component 1, which accounts for the greatest variation in the data, was driven by sex, additional sPLSDA was performed on males and females specifically to isolate any sex specific genotype effects. The top 10 loadings were shown for Component 1 (injury effect) and Component 2 (genotype effect) for sex-separated sPLSDA. For naïve mature mice, sPLSDA (5 components, 10 variables per component) was used to generate a scores plot that allowed for visualization of sample clustering by genotype. The top 10 loadings were shown for Component 1 (genotype effect). All sPLSDA were performed on auto scaled data (mean-centered and divided by the standard deviation of each variable).

Individual μCT parameter values are shown for contralateral limbs as well as ACLR limbs normalized to the respective sex- and genotype-matched contralateral to isolate genotype-dependent effects on injury induced bone remodeling. Individual μCT parameter values are also shown for the naïve, mature adult cohort.

*Paraffin histology and histopathological scoring*

### Suppl. Table 2. Synovitis histopathological scoring

| Synovitis severity scoring | | | | |
|---|---|---|---|---|
| **Category**<br>**Range:**<br>**Region(s):** | **0** | **1** | **2** | **3** |
| **Pannus**<br>**Range:** 0-3<br>**Region(s):**<br>1. Anterior synovium/tibia | None. | Mild: Pannus has migrated onto bone but is not encroaching on articular surface. It is in the transition zone where there is calcified cartilage, but you are not yet at the articular surface. | Moderate: Pannus has migrated through the transition zone and <1x cartilage depth onto the articular surface. | Severe: Pannus has migrated > 1x cartilage depth onto the articular surface. |
| **Bone erosion**<br>**Range:** 0-3<br>**Region(s):**<br>1. Anterior femur<br>2. Anterior tibia | None. | Partial thickness loss of cortical bone only. Wavy surface. | Focal complete loss of cortical bone - communication with marrow cavity at one small vascular communication site. A single large “offshoot” of cortical bone erosion. | Widespread complete loss of cortical bone - communication with marrow cavity at multiple sites or broad area loss of cortical bone. |
| **Synovial lining hyperplasia**<br>**Range:** 0-3<br>**Region(s):**<br>1. Anterior, superior synovium<br>2. Anterior, inferior synovium | 1 cell thick. | Mild: 2-3 cells thick. | Moderate: 4-5 cells thick. | Severe ≥6 cells thick. |
| **Subsynovial inflammation**<br>**Range:** 0-3<br>**Region(s):**<br>1. Anterior synovium | None. | One pocket of densely associated inflammatory cells. | Two to three pockets of densely associated inflammatory cells. These are focal areas of dense subsynovial WBC infiltrate – but still predominantly normal subsynovial areolar connective tissue present. | ≥ Four pockets of densely associated inflammatory cells. This is widespread dense subsynovial WBC infiltrate with markedly reduced or little/no normal areolar connective tissue evident and some lymphoid follicle formation. |
| **Synovial fibrosis**<br>**Range:** 0-3<br>**Region(s):**<br>1. Anterior synovium | None (less than 10% to account for the immediate sublining and regular matrix around blood vessels). | Dispersed fibrosis (10% to 1/3 of synovial area). | Moderate fibrosis (>1/3 to 2/3 of synovial area). | Severe fibrosis (>2/3 of synovial area). |
| **Synovial exudate**<br>**Range:** 0-1<br>**Region(s):**<br>1. Anterior synovium | None. | Infiltration of inflammatory cells (neutrophils, macrophages, and/or lymphocytes) or fibrin in the synovial cavity. | | |

### Suppl. Table 3. PTOA histopathological scoring

| PTOA severity scoring | | | | | | | | |
|---|---|---|---|---|---|---|---|---|
| **Category**<br>**Range:**<br>**Region(s):** | **0** | **1** | **2** | **3** | **4** | **5** | **6** | **7** |
| **Structural damage**<br>**Range:** 0-7<br>**Region(s):**<br>1. Femur<br>2. Tibia | Normal cartilage. | Roughened surface with small fibrillations, wavy articular surface. | Fibrillations immediately below superficial layer or some loss of laminal surface. | Horizontal cracks or separations between calcified and non-calcified cartilage. | Mild loss of non-calcified cartilage (<10% surface area). | Moderate loss of non-calcified cartilage (10-50% surface area). | Severe loss of non-calcified cartilage (>50% surface area). | Erosion of cartilage to subchondral bone (any percent surface area). |
| **Proteoglycan loss**<br>**Range:** 0-3<br>**Region(s):**<br>1. Femur<br>2. Tibia | Normal cartilage. | Decreased but not complete loss of Safranin-O staining in non-calcified areas. Saf-O loss does not penetrate completely from superficial to deep cartilage. | Saf-O loss penetrates completely from superficial to deep cartilage. Focal loss of Safranin-O staining in non-calcified area (<30% surface area). | Saf-O loss penetrates completely from superficial to deep cartilage. Diffuse loss of Safranin-O staining in non-calcified cartilage (>30% surface area). | | | | |
| **Chondrocyte hypertrophy**<br>**Range:** 0-1<br>**Region(s):**<br>1. Femur<br>2. Tibia | None. | Enlarged chondrocyte lacunae with lack of Saf-O stain around collapsed cell. | | | | | | |
| **Osteophyte size**<br>**Range:** 0-3<br>**Region(s):**<br>1. Femur<br>2. Tibia | None. | Small – ≤1x thickness as adjacent cartilage. | Medium – >1x to 3x as thick as adjacent cartilage. | Large - >3x thicker than adjacent cartilage. | | | | |
| **Osteophyte maturity**<br>**Range:** 0-3<br>**Region(s):**<br>1. Femur<br>2. Tibia | None. | Predominately cartilage. | Mixed cartilage and bone with vascular invasion. | Predominately bone. | | | | |
| **Subchondral bone thickening**<br>**Range:** 0-3<br>**Region(s):**<br>1. Femur<br>2. Tibia | Normal SCB. | Mild thickening, <50% increase. | Moderate thickening, 50-100% increase. | Severe thickening, >100% increase. | | | | |

*Bulk RNA sequencing*

Briefly, synovia were harvested into TRIzol, homogenized, and RNA was isolated using an RNeasy® Mini Kit (QIAGEN, Hilden, Germany). Samples were submitted to Novogene (San Francisco, California, United States) for sequencing and read count matrices were returned for bioinformatic analyses. Whole mRNA was sequenced using polyA capture (150 bp paired-end reads, >40M reads/sample) on a NovaSeq 6000 sequencer (Illumina Inc., San Diego, California, United States). Quality control of sequencing data confirmed high quality RNA (RIN > 7) and successful, accurate sequencing in all samples (**Suppl.-Tables 4-7**).

All bulk RNA-sequencing data analysis was performed in RStudio (R v4.4.0, RStudio v2024.04.1). Raw read counts were filtered to exclude non-protein-coding and sex-linked genes. Lowly-expressed genes were also removed, with only genes with FPKM > 1 and > 10 total reads in at least *n*=2-3 per group/genotype/sex samples retained. Outliers were assessed via a combination of principal component analysis (PCA) (**Suppl.-Fig. 11A**) and a skeletal muscle enrichment score (a gene module score based on GO:0003009 – Skeletal Muscle Enrichment) was calculated to assess skeletal muscle tissue contamination (**Suppl.-Fig. 11B**). One outlier was detected on PCA which exhibited significant skeletal muscle tissue contamination, concomitant with low expression of prominent synovial gene markers (e.g. synovial fibroblast marker, *Pdgfra*) – this outlier was removed from downstream analyses. PCA and Eigencor (PCAtools v2.14.0) analyses (**Suppl. Fig. 11C**) suggested that sex was not a driving factor of variance in the dataset – consequently, male and female samples were merged for downstream analyses to focus on conserved trends between sexes. Differential gene expression analysis was performed using weighted Limma-Voom[7] (limma v3.58.1) with injury and genotype as between-subject effects. DEGs were defined as $P_{adj} < 0.05$. Heatmaps and volcano plots illustrating gene-level findings were generated using *pheatmap* (v1.0.13) and *EnhancedVolcano* (v1.20.0) R packages, respectively[8,9]. Gene expression heatmaps are presented as Z-scores, normalized between conditions within each gene.

Pathway analyses were performed using PantherDB[10], as well as the *clusterProfiler*[11] (v4.10.1) implementation of fast gene set enrichment analysis (GSEA), both using the GO: Biological Processes database. Reactome GSEA was performed using the Reactome GSA R package. For GSEA, all genes (excluding sex-linked, non-protein coding, and non-expressed genes) were assigned signed gene rankings according to the formula sign(log2FC) * *P*-value (based on differential-expression analysis), sorted in descending order, and submitted to GSEA analysis via the *gseGO* function. GO terms were condensed to minimize redundancy by collapsing significant terms with a shared lineage into their parent term. For the GSEA using the GO:BP database and Reactome Pathway database, terms with $P_{adj} < .05$ were considered significant. A curated list of terms is shown and plotted by its normalized enrichment score (NES, positive indicates upregulation of a pathway in KO vs WT for that group, negative indicates downregulation). For PantherDB analysis, lists of significantly differentially-expressed genes ($P_{adj} < 0.05$) were submitted to statistical over-representation analysis (positively- and negatively-regulated genes were submitted separately). Full gene expression comparison lists, GSEA term lists, and PantherDB overrepresentation term lists can be found in **Supplemental Data File 1**.

*Statistical analysis:*

SPSS (v30, IBM, Armonk, NY) and Prism 9.0 (Graphpad, San Diego, CA) were used for statistical analyses. Multiple paired t-tests compared contralateral vs ACLR AngioSense 750 RID at 7d and 28d post-ACLR. T-test compared ACLR/Contralateral AngioSense 750 signal ratio across timepoints. Paired t-tests compared contralateral vs ACLR flow cytometry outputs and synovial *Thbs2* expression within each timepoint. T-tests compared *Thbs2* expression in *in vitro* synovial fibroblast treatments. For ProSense 680, AngioSense 750, histopathology, and μCT analyses in TSP2-KO and WT mice post-ACLR, two-way linear mixed-effects models with Sidak post-hoc correction compared continuous outcomes in the contralateral and ACLR groups separately (genotype and sex as between-subject factors). Raman spectroscopy analyses, two-way linear mixed-effects models with Sidak post-hoc correction compared continuous outcomes (genotype and sex as between-subject factors). In the ACLR and Raman spectroscopy studies, the effect of genotype within sex and the main effect of genotype across sexes was evaluated. T-tests compared bone morphometry metrics and histopathologic scoring in naïve, mature WT and TSP2 KO mice. Sparse partial least squares-discriminant analysis (sPLSDA) within MetaboAnalyst was used to perform dimensionality reduction of μCT data, and variable importance in projection (VIP) coefficients were calculated for the components describing sex-dependent, injury-dependent, and genotype-dependent variance.

### SUPPLEMENTAL FIGURE LEGENDS

**Supplemental Figure 1. A)** Top 10 upregulated and downregulated genes in mouse synovial endothelial cells when comparing Sham vs 7d ACLR and Sham vs 28d ACLR[2]. **B)** Individual ligand-receptor contributions to the THBS signaling pathway network in **Fig. 1G**.

**Supplemental Figure 2. A)** RT-qPCR validation of *Thbs2* gene expression upregulation in the synovium at 3d and 14d post-ACLR compared to the uninjured contralateral synovium (n=8-9 per timepoint; paired t-test between contralateral and ACLR synovia within each timepoint). **B)** Quantification of TSP2-GFP+ signal in cryosections of the contralateral and ACLR synovium at 14d and 28d post-ACLR (n=3 per timepoint; paired t-test between contralateral and ACLR synovia within each timepoint). **C)** Feature plots of mouse synovial single cell RNAseq data[2] from showing *Thbs2, Cd36,* and *Cd47* gene expression distribution across all cell types. **D)** Heatmaps of mouse synovial scRNAseq data showing expression of thrombospondin family genes and TSP2 receptor genes, *Cd36* and *Cd47*, within endothelial cells and fibroblasts[2]. **E)** qPCR plots of *Thbs2* expression in synovial fibroblasts after treatment with TNF-α, Wnt3a, or TGF-β (n=3-6 per group; t-test between each separate control and treatment group). All bars show mean ± SEM.

**Supplemental Figure 3. A)** The TSP2-GFP+ population was determined using a fluorescence-minus-one (FMO) control of synovial cells from a wild-type (WT) mouse. **B)** Gating strategy for flow cytometry. First, debris were excluded based on forward (FSC-A) and side (SSC-A) scatter profile (i), then doublets were excluded based on side scatter (SSC-H x SSC-W) (ii) and forward scatter (FSC-H x FSC-W) (iii). TOPRO3 was used to exclude dead cells (iv) then live TOPRO3- cells were plotted according to CD31 and CD45. Major cell types were defined as CD31- CD45- synovial fibroblasts (v), CD31+ endothelial cells (vi), or CD45+ hematopoietic cells (vii). CD45+ cells were then gated on and defined as either CD11b+ CD3- myeloid cells/macrophages (viii) or CD11b- CD3+ T cells (ix). **C)** Total number of TSP2-GFP+ CD31- CD45- cells in contralateral or 3d ACLR synovium. **D)** Percentage and **(E)** total number of TSP2-GFP+ CD45+ cells that were CD11b+ or CD11b- in contralateral or 3d ACLR. **F)** Median fluorescence intensity (MFI) of TSP2-GFP in live cells from contralateral or 3d ACLR synovium (n=4 mice). Paired two-tailed student's t-tests were performed to assess significance in (C-F). n.d.: not detected. All bars show mean ± SEM.

**Supplemental Figure 4. A)** ProSense 680 NIR signal intensity quantification of raw integrated density (RID) in contralateral limbs and the ACLR/Contralateral ProSense 680 RID signal ratio (n=5-6 per sex/genotype). **B)** AngioSense 750 NIR signal quantification expressed as RID in the 7d ACLR limb and contralateral limb with the ACLR/Contralateral AngioSense 750 RID signal ratio (n=6 per sex/genotype). All statistical analyses utilized a two-way linear mixed effects model to look at the effect of genotype within sex and the main effect of genotype across sex. All bars show mean ± SEM. **C)** Representative intravital fluorescence imaging heatmap of AngioSense 750 at 7d post-ACLR.

**Supplemental Figure 5.** Each of the individual subscores comprising the composite scores are shown. PTOA score is evaluated for the femur and tibia separately and includes structural damage, proteoglycan loss, chondrocyte hypertrophy, subchondral bone thickness, osteophyte size, osteophyte maturity. The total femoral and tibial PTOA score is a summation of the individual subscores for the femur and tibia, respectively, while the composite total joint PTOA score is the summation of the total femur and tibia scores. The synovitis score is a summation of the individual subscores evaluated in the anterior synovium including pannus, lining hyperplasia of the regions anterior/superior and anterior/inferior to the meniscus, fibrosis, subsynovial inflammation, exudate, femur bone erosion, and tibia bone erosion. These scores are shown for the **(A)** contralateral limb and **(B)** 7d ACLR limb. All statistical analyses utilized a two-way linear mixed effects model to look at the effect of genotype within sex and the main effect of genotype across sex. All bars show mean ± SEM.

**Supplemental Figure 6.** μCT analysis included evaluation of the metaphyseal and epiphyseal trabecular bone compartments along with the medial and lateral subchondral bone compartments. **A-D)** To isolate genotype driven differences in bony remodeling after joint injury, all raw μCT bone morphometry readouts were normalized to the appropriate sex- and genotype-matched contralateral limbs (n=8 per sex/genotype). **(E-H)** Individual μCT bone morphometry parameters are shown for contralateral limbs (n=8 per sex/genotype). All statistical analyses utilized a two-way linear mixed effects model to look at the effect of genotype within sex and the main effect of genotype across sex. All bars show mean ± SEM.

**Supplemental Figure 7. A-B)** Dimensionality reduction of female µCT bone morphometry data 28d post-ACLR. **A)** sPLSDA scores plot. **B)** Component 1 loadings driving injury-dependent separation (18.2% of data variation) and component 2 loadings driving genotype-dependent separation (19.5% of data variation) of the samples (n=8 per genotype). **C-D)** Dimensionality reduction of male µCT bone morphometry data 28d post-ACLR. **C)** sPLSDA scores plot. **D)** Component 1 loadings driving injury-dependent separation (26.6% of data variation) and component 2 loadings driving genotype-dependent separation (25.8% of data variation) of the samples (n=8 per genotype). Representative images of the anterior synovium in **(E)** contralateral and **(F)** 28d ACLR limbs. Representative images of the articular surfaces in **(G)** contralateral and **(H)** 28d ACLR joints.

**Supplemental Figure 8.** Each of the individual subscores comprising the composite scores are shown. PTOA score is evaluated for the femur and tibia separately and includes structural damage, proteoglycan loss, chondrocyte hypertrophy, subchondral bone thickness, osteophyte size, osteophyte maturity. The total femoral and tibial PTOA score is a summation of the individual subscores for the femur and tibia, respectively, while the composite total joint PTOA score is the summation of the total femur and tibia scores. The synovitis score is a summation of the individual subscores evaluated in the anterior synovium including pannus, lining hyperplasia of the regions anterior/superior and anterior/inferior to the meniscus, fibrosis, subsynovial inflammation, exudate, femur bone erosion, and tibia bone erosion. These scores are shown for the **(A)** contralateral limb and **(B)** 28d ACLR limb. All statistical analyses utilized a two-way linear mixed effects model to look at the effect of genotype within sex and the main effect of genotype across sex. All bars show mean ± SEM.

**Supplemental Figure 9.** Raman spectroscopy analysis of (**A**) subchondral bone and (**B**) epiphyseal trabecular bone in naïve mice (n=2 per sex/genotype with n=3 sites averaged together within each bone compartment per mouse). All Raman ratios were calculated from peak areas. All statistical analyses utilized a two-way linear mixed effects model to look at the effect of genotype within sex and the main effect of genotype across sex. All bars show mean ± SEM.

**Supplemental Figure 10. A-D)** μCT analysis evaluated the metaphyseal and epiphyseal trabecular bone compartments along with the medial and lateral subchondral bone compartments. Individual μCT bone morphometry parameters are shown (n=8 per genotype). **E)** sPLSDA scores plot with 95% confidence intervals shown. **F)** Component 1 loadings driving genotype-dependent separation (31.1% of variation). **G)** Representative images of Safranin-Orange and Fast Green-stained joints. **(H)** Each of the subscores comprising the composite scores are shown. OA score is evaluated for the femur and tibia separately and includes structural damage, proteoglycan loss, chondrocyte hypertrophy, subchondral bone thickness, osteophyte size, osteophyte maturity. The total femoral and tibial OA score is a summation of the individual subscores while the composite total joint OA score is the summation of the total femur and tibia scores. The synovitis score is a summation of the individual subscores evaluated in the anterior synovium including pannus, lining hyperplasia of the regions anterior/superior and anterior/inferior to the meniscus, fibrosis, subsynovial inflammation, exudate, femur bone erosion, and tibia bone erosion (n=11 per genotype). All statistical analyses utilized a t-test to look at the effect of genotype. All bars show mean ± SEM.

**Supplemental Figure 11. A-B)** Principal component analysis (PCA) and muscle gene set enrichment score were utilized to identify outliers in the data (n=3 per sex/group/genotype). One outlier was identified, sample KO_FS_3. **A)** PCA plot of all samples with 95% confidence interval with either group (Sham, 7d ACLR, 28d ACLR) or genotype (WT, KO) indicated. **B)** Muscle gene set enrichment score, based on variance-stabilized transformation (VST)-normalized expression. Each individual sample is shown. **C)** Eigencore plot showing the correlation coefficient of each factor for each of the top ten principal components.

**Supplemental Figure 12. Bulk RNAseq of TSP2 KO and WT synovium supplemental analysis. A)** Volcano plot showing DEGs comparing WT and KO synovia within the 7d ACLR and 28d ACLR groups. **B)** Heatmaps showing all genes with each condition. **C)** Gene set enrichment analysis comparing KO vs WT by group using the Reactome Pathway database. **D)** PantherDB gene ontology analysis of shared DEGs between Sham KO vs Sham WT and WT 7d ACLR vs WT Sham (column #1) and Sham KO vs Sham WT and WT 28d ACLR vs WT Sham (column #2). A curated list of terms shown is shown.
